# CtIP -mediated alternative mRNA splicing finetunes the DNA damage response

**DOI:** 10.1101/849547

**Authors:** Rosario Prados-Carvajal, Guillermo Rodríguez-Real, Gabriel Gutierrez-Pozo, Pablo Huertas

**Affiliations:** Departamento de Genética, Universidad de Sevilla, Sevilla, 41080, Spain; Centro Andaluz de Biología Molecular y Medicina Regenerativa-CABIMER, Universidad de Sevilla-CSIC-Universidad Pablo de Olavide, Sevilla, 41092, Spain

**Keywords:** CtIP, SF3B complex, PIF1, DNA damage response, mRNA splicing

## Abstract

In order to survive to the exposure of DNA damaging agents, cells activate a complex response that coordinates the cellular metabolism, cell cycle progression and DNA repair. Among many other events, recent evidence has described global changes in mRNA splicing in cells treated with genotoxic agents. Here, we explore further this DNA damage-dependent alternative splicing. Indeed, we show that both the splicing factor SF3B2 and the repair protein CtIP contribute to the global pattern of splicing both in cells treated or not to DNA damaging agents. Additionally, we focus on a specific DNA damage- and CtIP-dependent alternative splicing event of the helicase PIF1 and explore its relevance for the survival of cells upon exposure to ionizing radiation. Indeed, we described how the nuclear, active form of PIF1 is substituted by a splicing variant, named vPIF1, in a fashion that requires both the presence of DNA damage and CtIP. Interestingly, timely expression of vPIF1 is required for optimal survival to exposure to DNA damaging agents, but early expression of this isoform delays early events of the DNA damage response. On the contrary, expression of the full length PIF1 facilitates those early events, but increases the sensitivity to DNA damaging agents if the expression is maintained long-term.

## INTRODUCTION

DNA is constantly threatened by endogenous and external sources that compromise its integrity. Thus, during evolution eukaryotes have developed a complex signaling network that finetunes the response to those threats. Generally referred as the DNA Damage Response (DDR), such network affects virtually every aspect of the cell metabolism (Ciccia and Elledge, 2010; Jackson and Bartek, 2009). In addition to those changes, the DDR activates the actual repair of damaged DNA. There are many different DNA lesions, thus several specific repair pathways coexist. DNA double strand breaks (DSBs) are the most cytotoxic form of DNA damage. Indeed, repair of DSBs can be achieved by different mechanisms, generally grouped in two categories, regarding the use or not of a template for repair. Whereas non-homologous end-joining (NHEJ) uses no homology to seal DSBs, homologous recombination will copy the information from a homologous sequence (Jasin and Rothstein, 2013; Lieber, 2010). The decision between those pathways is controlled by the DDR, and relies on the activation or not of the processing of the DNA ends, the so-called DNA end resection (Symington et al., 2014). This regulation is mostly achieved by controlling a single protein, CtIP, which integrates multiple signals in order to activate or not end processing (Makharashvili and Paull, 2015; Symington et al., 2014). Thus, in order to modulate resection and, as a consequence, homologous recombination, CtIP works together with several other proteins that affect the processivity of the end resection, mainly the tumor suppressor gene BRCA1 (Cruz-García et al., 2014).

Recently, a crosstalk between the DDR and the RNA metabolism at different levels has been discovered. Indeed, the number of factors that participate in the DNA damage response and/or are regulated by it has expanded considerably in recent years to include many RNA-related proteins, notably splicing and alternative splicing factors (Jimeno et al., 2019; Nimeth et al., 2020). So, post-translational changes of splicing factors following DNA damage such as phosphorylation, ubiquitination, sumoylation, neddylation, PARylation, acetylation and methylation of splicing factors, have been documented. On the other hand, bona fide DDR factors also directly control splicing. For example, BRCA1 regulates alternative splicing in response to DSB formation through its DNA damage-dependent interaction with several splicing factors such as SF3B1, one of the subunits of the Splicing Factor 3B (SF3B) complex (Savage et al., 2014). SF3B is a multiprotein complex essential for the accurate excision of introns from pre-messenger RNA (Golas et al., 2003). This complex consists of seven subunits: SF3B1 (also known as SF3b155), SF3B2 (SF3b145), SF3B3 (SF3b130), SF3B4 (SF3b49), SF3B5 (SF3b10), SF3B6 (SF314a) and SF3B7 (PHF5a) (Spadaccini et al., 2006). SF3B plays an indispensable role during the assembly of the pre-spliceosome recognizing the intron’s branch point (Teng et al., 2017). Interestingly, several subunits of this complex have been found in genome wide screens for factors involved in DNA repair, affecting homologous recombination (Adamson et al., 2012), controlling genome stability (Paulsen et al., 2009) or as substrates of the checkpoint kinases (Matsuoka et al., 2007). Moreover, we recently reported that the SF3B complex directly interacts with CtIP and regulates its activity in DNA end resection (Prados-Carvajal et al., 2018).

Interestingly, in addition to its well defined role in DSB repair by regulating DNA end resection, CtIP seems to perform many additional tasks in the cell, affecting DNA repair, cell cycle progression, checkpoint activation, replication and transcription (Duquette et al., 2012; Liu and Lee, 2006; Makharashvili and Paull, 2015; Moiola et al., 2012; Wu and Lee, 2006). CtIP promotes the expression of several genes, such as Cyclin D1, and also activates its own promoter (Liu and Lee, 2006). The role of CtIP in regulating gene expression is confirmed by its interaction with other transcriptional factors like IIKZF1, TRIB3 and LMO4 (Koipally and Georgopoulos, 2002; Sum et al., 2002; Xu et al., 2007). Also, CtIP contributes to DNA damage-dependent cell cycle arrest in S and G2 phases promoting p21 transcription (Li et al., 1999; Liu et al., 2014) and upregulating the expression of GADD45A (Liao et al., 2010). Additionally, CtIP has been reported to regulate R-loop biology. CtIP deficiency has been shown to promote the accumulation of stalled RNA polymerase and DNA:RNA hybrids at sites of highly expressed genes (Makharashvili et al., 2018). On the contrary, CtIP depletion reduces DNA:RNA hybrid accumulation dependent on *de novo* transcription of dilncRNA (damage-induced long non-coding RNAs) starting at DSBs (D’Alessandro et al., 2018). Hence, CtIP loss seems to increase R-loops that are produced as a consequence of previous transcription and appears to decrease *de novo* production of diRNAs (DSB-induced small RNA), thus reducing the DNA:RNA hybrids formed after DNA damage.

Hence, CtIP has a central role in the DDR and DNA repair, but plays additional roles in RNA biology. As mentioned before, it physically interacts with the SF3B splicing complex (Prados-Carvajal et al., 2018). Thus, we wondered whether CtIP-SF3B functional relationship might extend to controlling mRNA splicing and, more specifically, DNA damage-induced alternative splicing. Here we show that both SF3B and CtIP, albeit in a more modest manner, influence expression and splicing of hundreds of genes. This effect is visible in unchallenged cells, but more evident when cells have been exposed to a DNA damaging agent. Then, we analyzed in detail the effect of a DNA damage- and CtIP-dependent alternative splicing event of the helicase PIF1. Although PIF1 and CtIP also interact directly and are involved in DNA end resection (Jimeno et al., 2018a), we observed that such alternative splicing of PIF1 is not involved in DNA end processing but it affects the cell survival upon exposure to DNA damaging agents. Interestingly, this alternative PIF1 form, when expressed constitutively, hampers the recruitment of DSB repair proteins at early time points but makes cells hyper-resistant to treatments with camptothecin.

## RESULTS

### SF3B controls DNA damage-induced alternative splicing

As mentioned in the introduction, SF3B controls HR and DNA end resection (Prados-Carvajal et al., 2018). Whereas the resection phenotype was completely dependent on regulation of CtIP, our data suggested other, CtIP-independent, roles of SF3B in DNA repair (Prados-Carvajal et al., 2018). Due to the well stablished role of SF3B in splicing (Sun, 2020) and, particularly its implication in DNA damage-dependent alternative splicing (Savage et al., 2014), we decided to analyze this role in more detail. Thus, we carried out a splicing microarray Transcriptome Arrays HTA & MTA using both damaged (6 hours after 10 Gy of irradiation) or untreated cells that were depleted or not for SF3B2 using shRNA (Figure 1A; see Materials and Methods for details). As previously published, SF3B2 depletion affects CtIP protein levels slightly (Prados-Carvajal et al., 2018). Such array allows the genome-wide study of RNA expression and RNA splicing simultaneously. As SF3B2 controls the levels of CtIP and BRCA1 mRNA (Prados-Carvajal et al., 2018), we first focused on total RNA levels genome-wide. Changes were considered significant when the fold change (FC) was 2 or more and the p-value less than 0.05. Indeed, SF3B2 depletion using shRNA leads to the specific upregulation of 52 genes solely in undamaged conditions when compared with control cells (Figure 1B). Moreover, 26 genes were exclusively overexpressed in SF3B2 downregulated cells upon exposure to DNA damage (Figure 1B). Additionally, mRNAs abundance from 27 genes was increased in cells with reduced SF3B2 levels in both damaged and undamaged cells (Figure 1B). A list of those genes could be found in Table 1. On the other hand, SF3B2 depletion also reduced expression of 97 genes (Figure 1C and Table 2). Only 10 of those genes were downregulated specifically in unperturbed conditions, 82 in cells that were exposed to ionizing radiation and 5 in both conditions, including SF3B2 itself as expected due to the shRNA-induced downregulation (Figure 1C, Table 2).

**Table 1:** Genes upregulated upon SF3B2 depletion.

| **Genes upregulated in undamaged cells** | **Genes upregulated in damaged cells** | **Genes upregulated in undamaged and damaged cells** |
| --- | --- | --- |
| ABCA5 | ANKRD18DP | BET1 |
| AC087073.1 | ANKRD45 | C12orf39 |
| ANTXR2 | CCDC30 | CBWD1 |
| ATG12 | CHKA | CBWD2 |
| C14orf37 | CTD-2651B20.3 | CBWD3 |
| C1orf168 | EIF5 | CBWD6 |
| C1orf63 | GABRR3 | CBWD7 |
| C21orf91 | HAVCR2 | CDO1 |
| C9orf156 | JAKMIP1 | CGRRF1 |
| CCDC66 | KLHL41 | CTD-2023N9.3 |
| CD274 | NME8 | FAM133B |
| CDKN1B | PDE4D | FLVCR1 |
| CREBRF | RP11-386I8.6 | GMFB |
| DNAJC27 | RP11-473L15.3 | KDM3A |
| DTD2 | RP11-545I10.2 | MITD1 |
| ENOX1 | RP4-553F4.6 | MMP13 |
| FBXW7 | RRH | MTHFD2L |
| GBP1 | SLFN12 | NPM3 |
| GDAP1 | SPATA9 | NQO1 |
| GTF2E2 | SYNE1 | RP11-111F5.3 |
| HEBP1 | THEMIS | RSAD2 |
| HSD17B13 | TRAM1 | SPHAR |
| IKZF3 | TSPY5P | SRD5A3-AS1 |
| IL18R1 | WDR87 | STARD4 |
| IL5 | ZNF442 | TULP3 |
| ITPR2 | ZNF571 | TXNDC2 |
| KANSL2 |  | ZNF844 |
| KIAA1467 |  |  |
| MAGOH |  |  |
| METTL13 |  |  |
| NCAPG2 |  |  |
| PARP16 |  |  |
| PCNXL2 |  |  |
| PDIK1L |  |  |
| PPARGC1A |  |  |
| RP11-159G9.5 |  |  |
| RP11-213G2.2 |  |  |
| RP11-261C10.3 |  |  |
| RP11-46C20.1 |  |  |
| RP13-996F3.4 |  |  |
| S100A8 |  |  |
| SCO1 |  |  |
| SESN1 |  |  |
| SLC25A27 |  |  |
| SRPRB |  |  |
| SUN3 |  |  |
| THOC7 |  |  |
| TP53INP1 |  |  |
| TRPM7 |  |  |
| USP54 |  |  |
| ZMAT3 |  |  |
| ZNF669 |  |  |

**Table 2:**
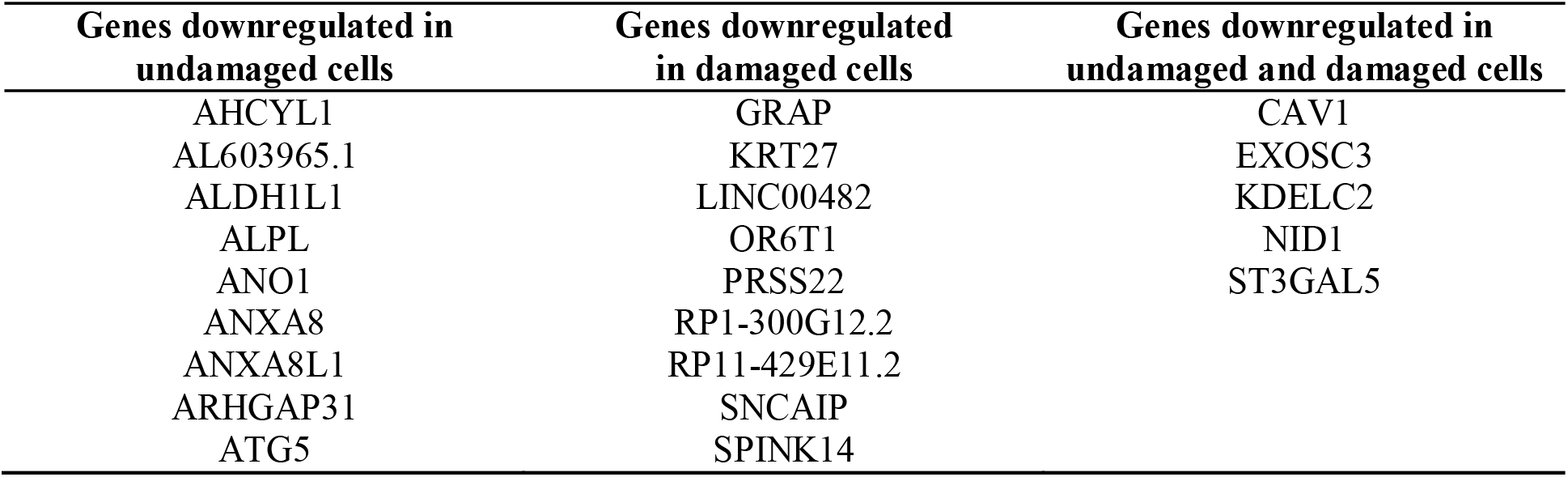
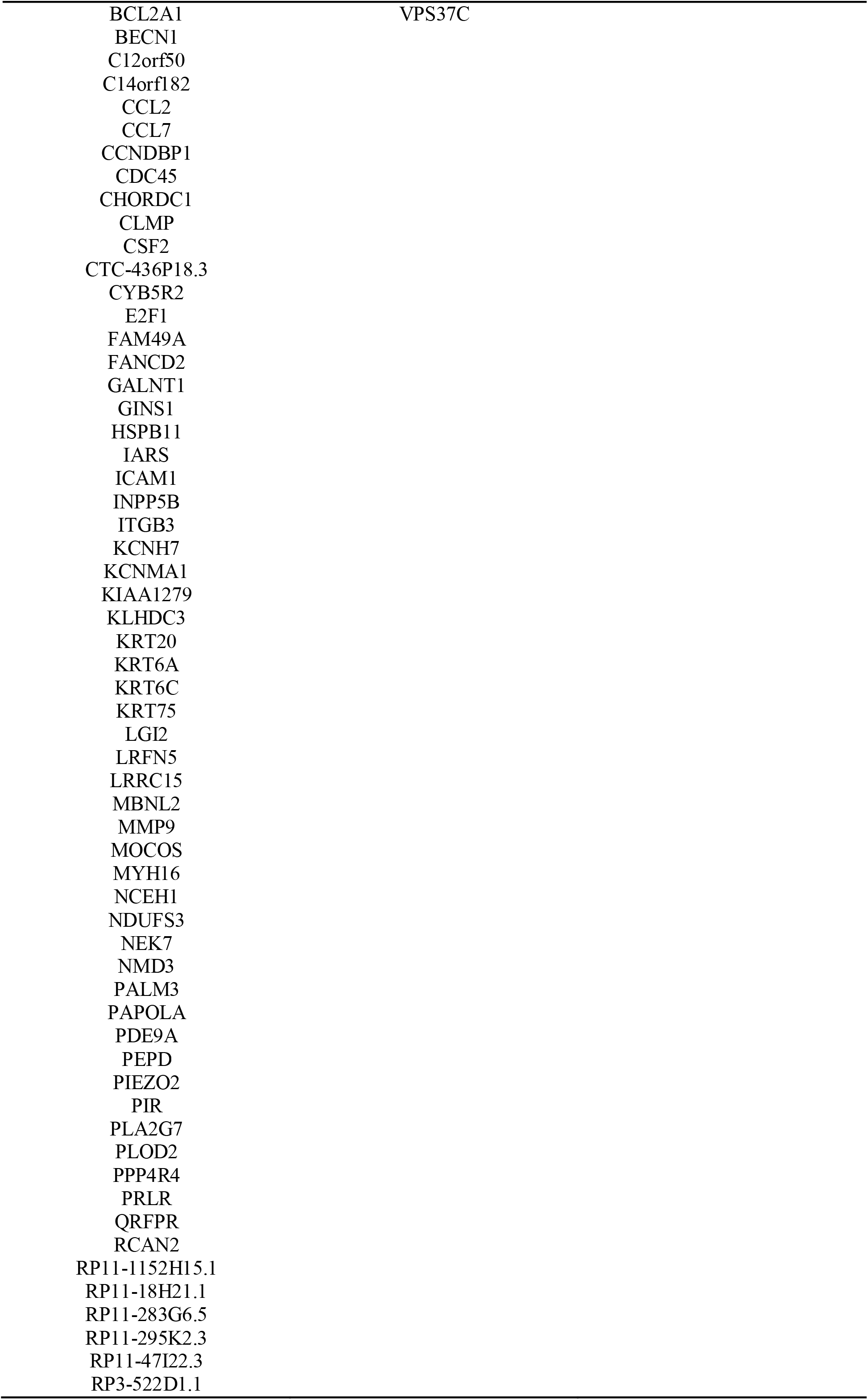
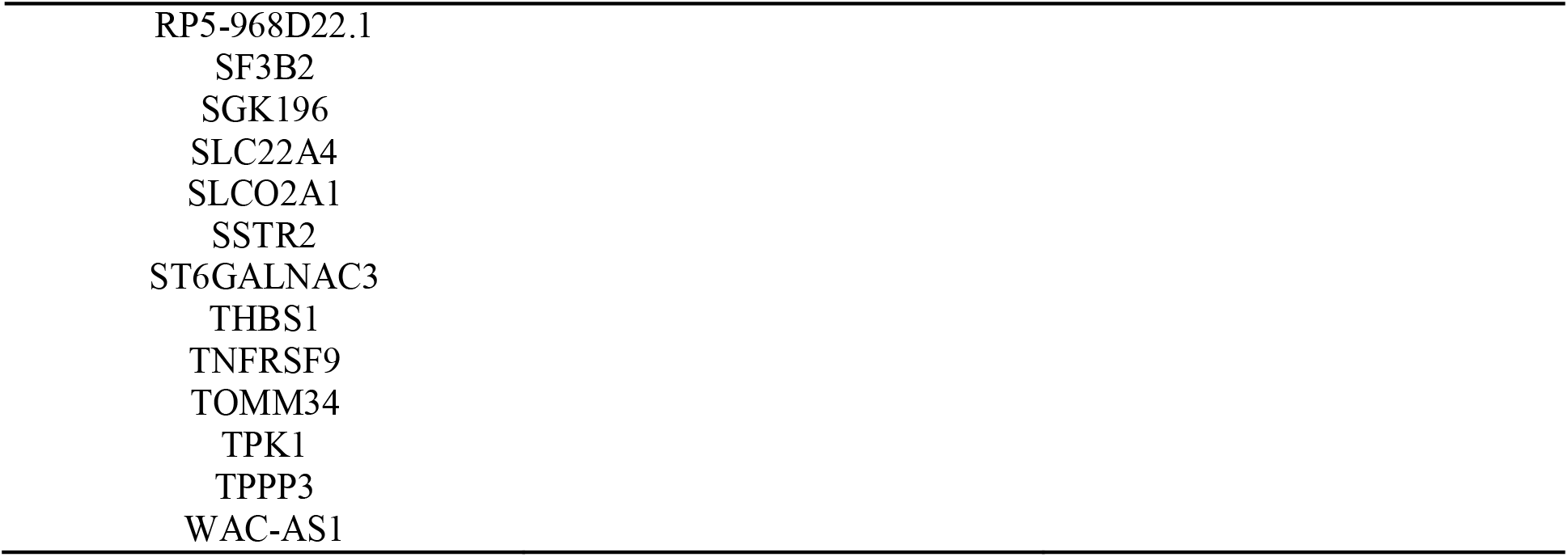
Genes downregulated upon SF3B2 depletion.

**Figure 1:**
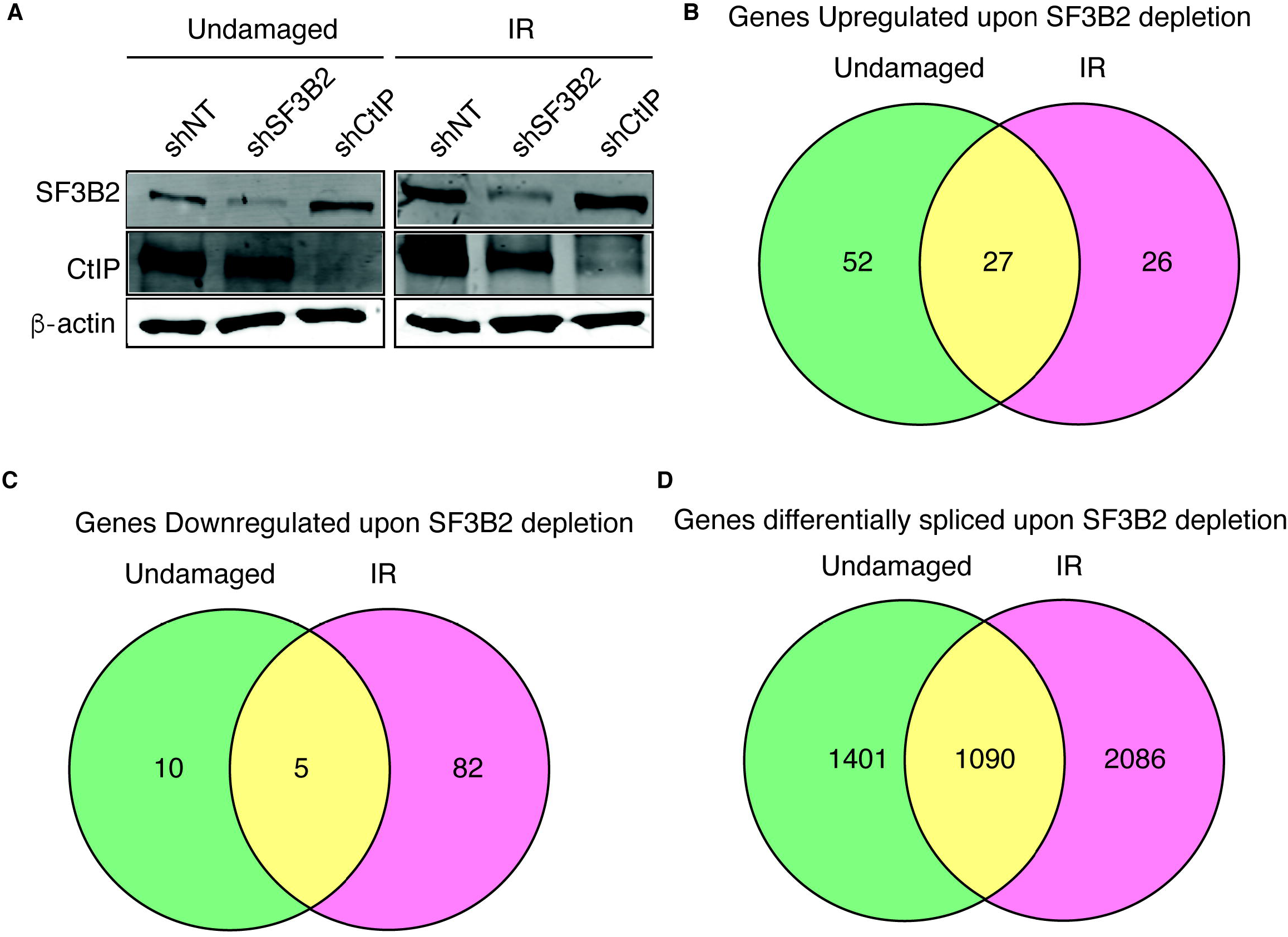
SF3B2 depletion affects gene expression and splicing of many genes. **A**, Representative western blot showing the expression levels of SF3B2 and CtIP upon depletion with the indicated shRNAs in cells exposed or not to 10 Gy of ionizing radiation. α-tubulin blot was used as loading control. **B**, Distribution of the genes upregulated upon SF3B2 depletion regarding the exposure or not to DNA damage. Fold change (FC)>2, p-value<0.05. Gene expression was measured using the GeneChip HTA Array as described in the methods section. The number of upregulated genes in undamaged cells (green), 6 h after exposure to irradiation (10 Gy; pink) or both (yellow) is shown in a Venn diagram. The actual list of genes can be found in Table 1. **C**, Same as A but for downregulated genes. The actual list of genes can be found in Table 2. **D**, the GeneChip HTA Array was used to look for genes that changed their splicing upon SF3B2 depletion, as mentioned in the methods section. Other details as in A. The actual list of genes can be found in Supplementary table 1.

In terms of mRNA splicing, we confirmed that both depletion of SF3B2 with shRNA and DNA damage induction affect such RNA processing globally. We first calculated the splicing index of exons on all four conditions and compared them *in silico* (FC>2, p-value<0.05; see Materials and Methods for additional details). The results are summarized in Figure 1C and a list of genes can be found in Supplementary Table S1. More than 4500 genes were differentially spliced when SF3B2 was absent, compared with a non-targeted shRNA (Figure 1D). Almost 25% did so regardless of the presence or absence of an exogenous source of DNA damage (yellow), but almost 45% showed splicing events that were both DNA damage- and SF3B-dependent (red) and only 30% of the genes were spliced by SF3B in undamaged conditions (green). Thus, most of the splicing events that require SF3B2 happens in damaged samples, indicating that this factor is especially relevant in stress conditions.

Indeed, a different analysis considering all genes that show an alternative splicing upon irradiation (IR) indicates that only 14% did so both in control and in SF3B2 depleted cells, whereas 46% of the genes suffer DNA damage-induced alternative splicing only when SF3B2 was present, suggesting they require this factor for such event. Strikingly, an additional 40% of the genes suffer damage induced alternative splicing specifically in SF3B2 depleted cells, indicating that when the SF3B complex was absent the splicing landscape is severely affected and new events appear.

Interestingly, the pattern of gain (+) or loss (-) of specific events of alternative splicing was similar in all situations (Table 3) with the exception of SF3B2-depleted cells upon irradiation, that was more pronounced in agreement with a strong role of the SF3B complex in DNA damage-induced alternative splicing (Savage et al., 2014). Despite the strong quantitative difference in splicing, qualitatively the types of events were similarly distributed in all cases (Table 3).

**Table 3:** Specific splicing changes in response to IR and SF3B2 depletion. (+) represents gain and (-) loss of each specific event

| Comparison | Irradiated versus non irradiated samples in control cells | Irradiated versus non irradiated samples in SF3B2 depleted cells | SF3B2 depleted versus control cells in untreated conditions | SF3B2 depleted versus control cells in irradiated conditions |
| --- | --- | --- | --- | --- |
| (+) alt-C-terminus | 274 | 220 | 201 | 375 |
| (+) alt-N-terminus | 265 | 215 | 185 | 335 |
| (+) alt-coding | 39 | 44 | 35 | 53 |
| (+) nonsense mediated decay | 72 | 53 | 59 | 117 |
| (+) retained intron | 75 | 67 | 62 | 122 |
| (+) truncated | 43 | 52 | 54 | 115 |
| (-) alt-C-terminus | 228 | 234 | 231 | 503 |
| (-) alt-N-terminus | 245 | 249 | 222 | 479 |
| (-) alt-coding | 42 | 53 | 93 | 164 |
| (-) nonsense mediated decay | 44 | 44 | 77 | 152 |
| (-) retained intron | 61 | 68 | 76 | 175 |
| (-) truncated | 63 | 56 | 28 | 66 |

In order to validate the array, we studied the mRNA level of several splicing variants of genes that were identified as positives in the analysis. Due to our interest, we focused mainly on those that are related to DNA resection, recombination or the DNA damage response. To do so, we depleted SF3B2 using siRNA and induced or not DNA damage (10 Gy irradiation; Figure 2A). Cells were incubated for 6 hours to allow accumulation of DNA damage-dependent isoforms. We used qPCR and sets of isoform-specific primers (Table 4) to study the level of different variants. In all cases, we included an analysis of a “common isoform” that is present ubiquitously in all conditions. The ratio between the alternative variant and the “common isoform” was normalized to control cells, i.e. non-irradiated cells transfected with non-targeted siRNA. As shown in Figure 2B, the levels of a specific *BRCA1* isoform increased upon DNA damage, but SF3B2 depletion blocks the accumulation of such *BRCA1* mRNA specie even in untreated cells. Thus, we described a SF3B2- and DNA-damage induced splicing variant of this mRNA. Differently, we confirmed that *RAD51* and *EXO1* have a damaged-dependent isoform that is independent of SF3B2 (Figures 2C and 2D). Also, in agreement with the array data, the levels of a *DNA2* mRNA variant increased specifically with DNA damage in the absence of SF3B2 (Figure 2E). *PIF1* mRNA alternative isoform expression increased both upon irradiation and upon SF3B2 depletion in an epistatic manner (Figure 2F). Finally, we studied *ATR*, whose alternative splicing is dependent on SF3B2 regardless the presence or absence of DNA damage (Figure 2G). In summary, in agreement with the array, SF3B complex and/or DNA damage presence controls the alternative splicing of DNA repair factors.

**Table 4:**
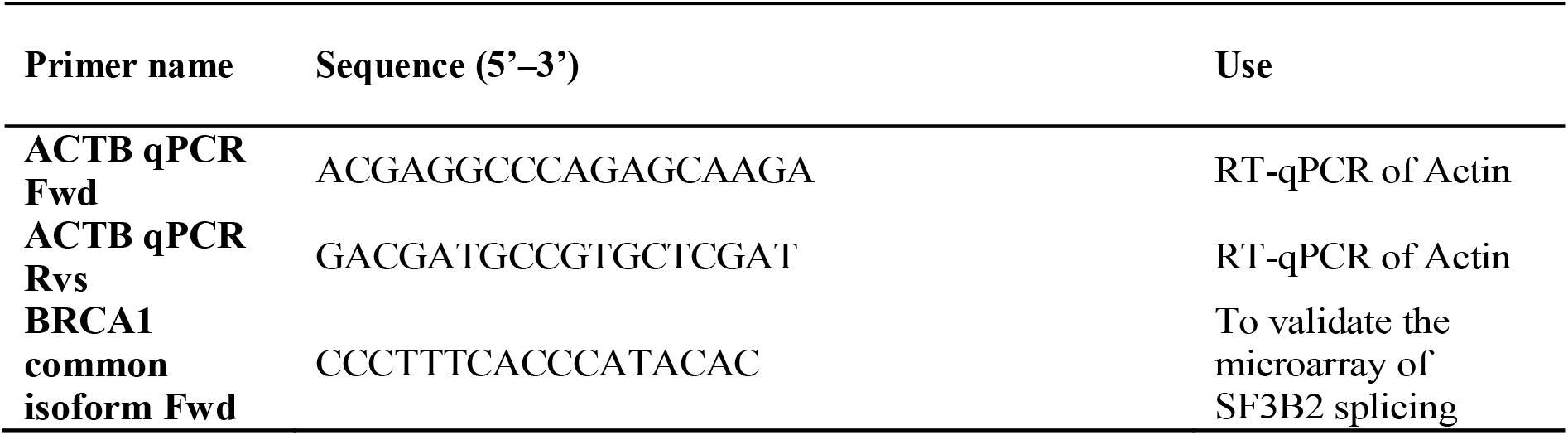
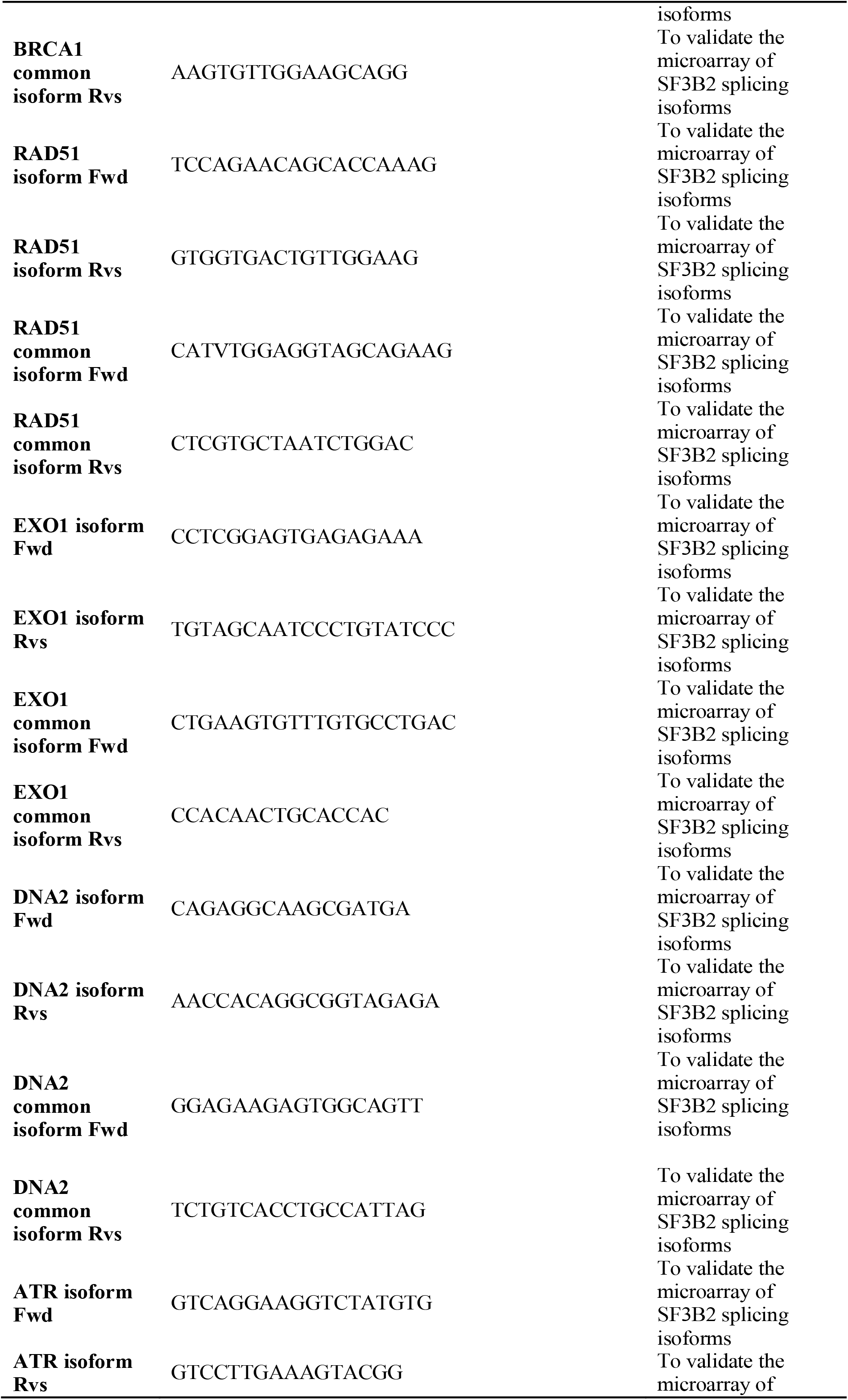
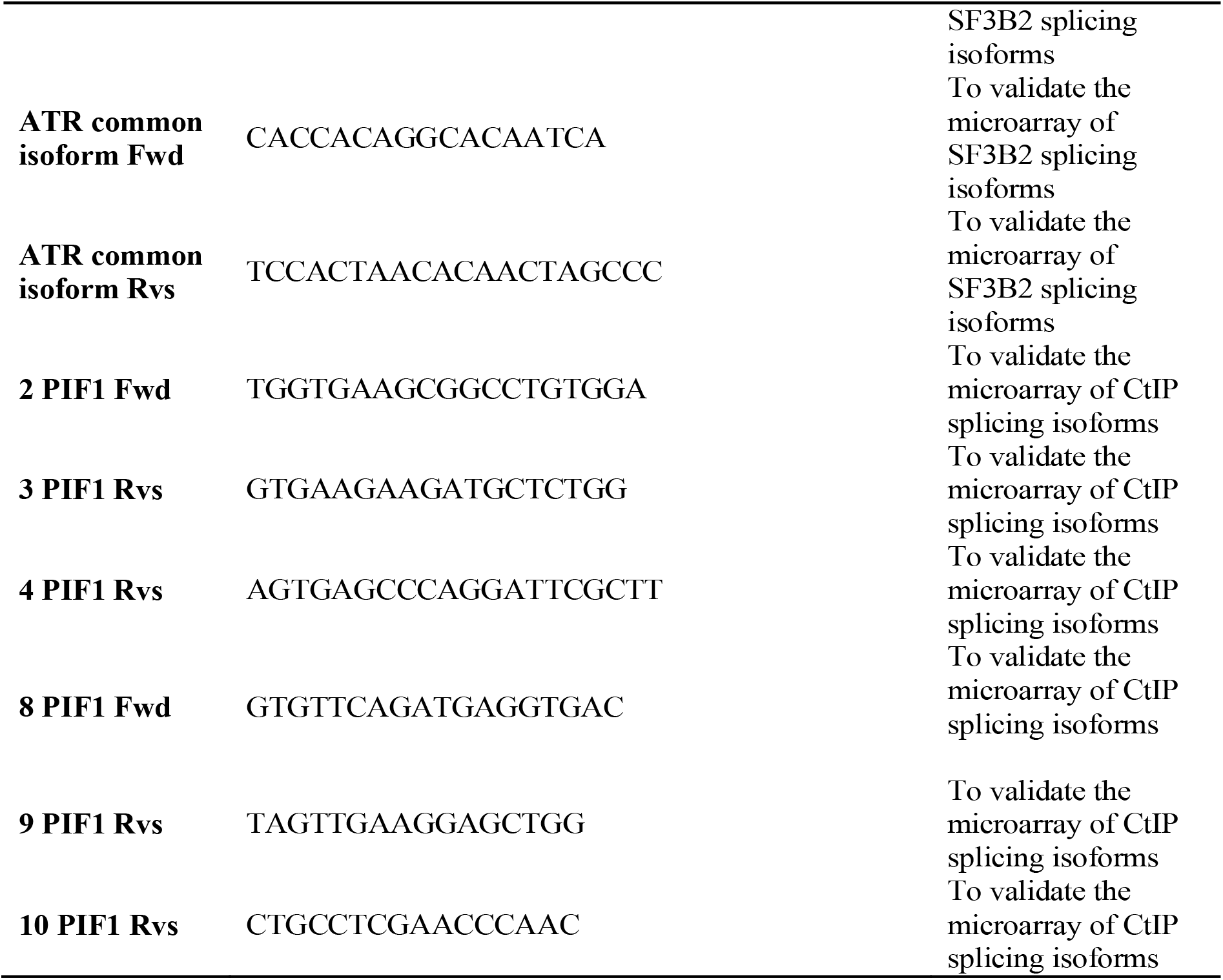
DNA primers.

**Figure 2:**
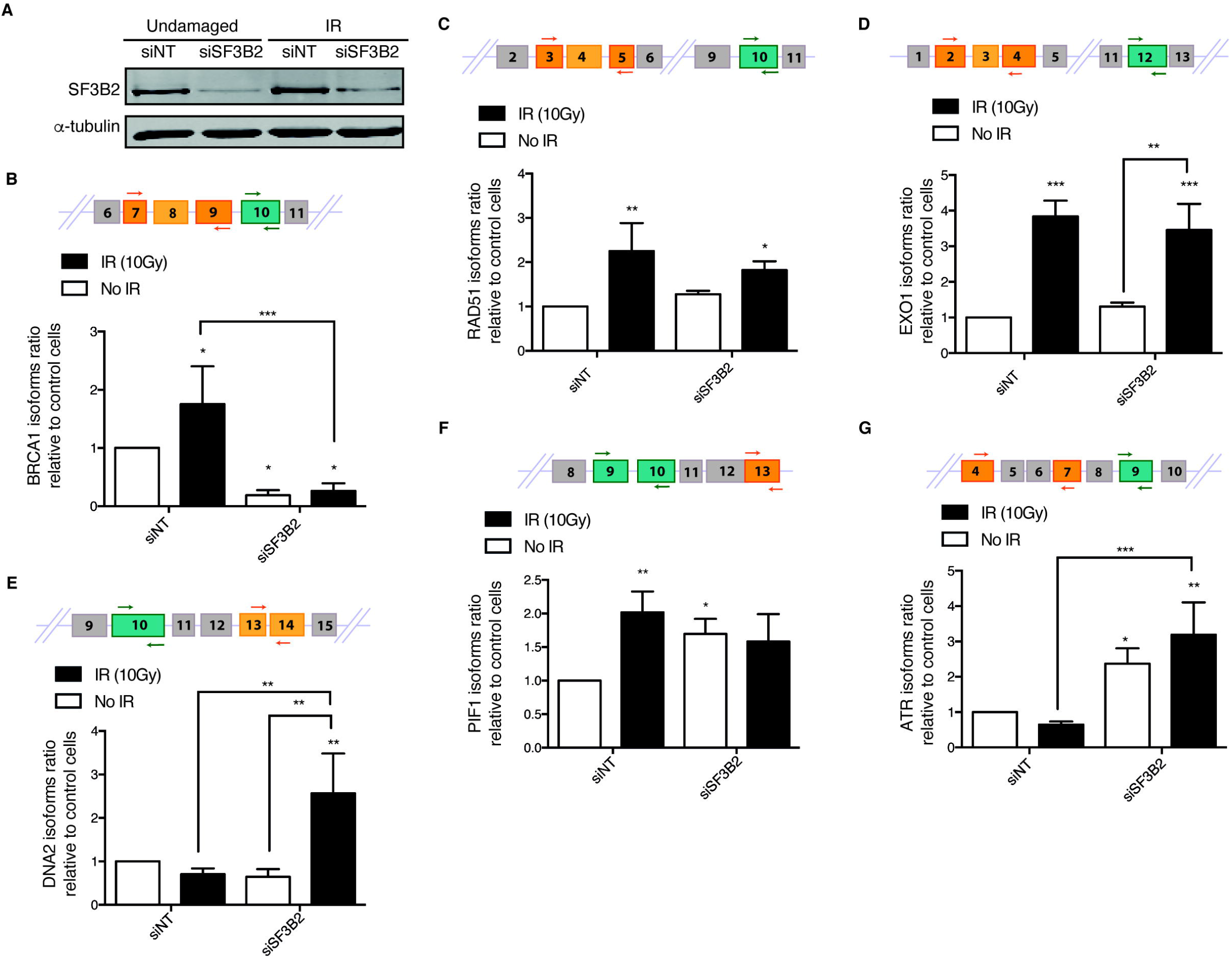
Splicing changes in DDR factors in cells depleted for SF3B2. **A**, Representative western blot showing the expression levels of SF3B2 upon depletion with a siRNA against SF3B2 or a control sequence (siNT) in cells exposed or not to 10 Gy of ionizing radiation. α-tubulin blot was used as loading control. **B-G**, Specific RNA isoforms levels of the indicated genes were calculated as the ratio between the abundance of the specific splicing form normalized with the total amount of each gene RNA by quantitative RT-PCR using specific primers in cells transfected with the indicated siRNAs and 6 hours after irradiation or mock treatment. See Materials and Methods for details. A schematic representation of the splicing events measured is shown in each case on the top. The common splicing event analyzed is shown in green, oligos are represented as arrows. The specific splicing that changes upon SF3B2 depletion is shown in orange. The graphs represent the average and standard deviation of three independent experiments. Statistical significance was calculated using an ANOVA test. * p<0.05,** p<0.01 and *** p<0.005.

### CtIP controls mRNA expression and splicing of several genes

As mentioned previously, SF3B directly interacts and regulates the resection factor CtIP. Interestingly, CtIP is a multifunctional protein that works in DNA repair, but also in other processes, including transcription (Wu and Lee, 2006). Moreover, other proteins related to DNA end resection, such as BRCA1, also have a role in RNA metabolism (Kleiman et al., 2005; Veras et al., 2009). Thus, we wondered whether CtIP could also play role in RNA splicing due to its connection with SF3B. To test this idea, and as for SF3B2, we used the splicing microarray to analyze RNA abundance and splicing genome-wide in cells depleted or not for CtIP using an shRNA, both in damaged and untreated conditions (Figure 1A for depletion of CtIP). When studying genome wide expression level of human genes, we observed that upon depletion of CtIP, and despite the assigned function in transcription, only 74 mRNAs showed altered abundance: 36 were upregulated and 38 downregulated (Figure 3A and B and Tables 5 and 6; Note that CtIP itself is among the downregulated ones, as expected due to the effect of the shRNA). Only 12 genes were exclusively upregulated in cells exposed to IR in cells depleted for CtIP compared with control cells (Figure 3A and Table 5). However, in undamaged conditions solely 22 genes were upregulated and the levels of only two genes increased in both conditions, with and without damage, in cells downregulated for CtIP (Figure 3A and Table 5). On the other hand, CtIP knockdown reduced the expression of 16 genes exclusively in unperturbed cells whereas the expression of 18 genes were decreased in irradiated cells (Figure 3B and Table 6). The expression of only 4 genes was downregulated upon CtIP depletion in both damaged and undamaged cells (Figure 3B and Table 6).

**Table 5:** Genes upregulated upon CtIP depletion.

| Genes upregulated in undamaged cells | Genes upregulated in damaged cells | Genes upregulated in undamaged and damaged cells |
| --- | --- | --- |
| AC010967.2 | AC068535.3 | MMP13 |
| AC016582.2 | ADAMTS9 | TULP3 |
| AC064852.4 | CD200R1L |  |
| AC073043.2 | DMXL2 |  |
| AF064858.7 | IL10 |  |
| ARL14 | IL24 |  |
| C9orf153 | KLHL6 |  |
| C9orf156 | MMP3 |  |
| CBLN2 | RP11-445K13.2 |  |
| COL10A1 | SPATA9 |  |
| EGLN1 | SRD5A3-AS1 |  |
| IKZF3 | WDR87 |  |
| KRT13 |  |  |
| NCEH1 |  |  |
| PARP16 |  |  |
| PPP2R1B |  |  |
| PRSS44 |  |  |
| RP11-285B24.1 |  |  |
| RP11-290K4.1 |  |  |
| RP11-300J18.2 |  |  |
| RP11-9G1.3 |  |  |
| RP4-553F4.6 |  |  |
| RP4-650F12.2 |  |  |
| SGIP1 |  |  |
| U51244.2 |  |  |

**Table 6:** Genes downregulated upon CtIP depletion.

| Genes downregulated in undamaged cells | Genes downregulated in damaged cells | Genes downregulated in undamaged and damaged cells |
| --- | --- | --- |
| DACT3 | C8orf4 | CtIP |
| DRAP1 | CCL7 | GALNT7 |
| ELOVL7 | CD69 | OLR1 |
| FGF9 | CTGF | SMYD2 |
| LINC00482 | CXCL10 |  |
| MYOCD | GOLT1A |  |
| NDP | INPP4B |  |
| OR6T1 | KAT2B |  |
| RBMS3-AS1 | MKX |  |
| RCN2 | NINJ1 |  |
| RP11-1069G10.1 | NTRK3 |  |
| SEMA3D | PDCD4 |  |
| SEZ6 | RP5-1172A22.1 |  |
| SMIM12-AS1 | RP5-968D22.1 |  |
| SNORA38B | S1PR3 |  |
| VAV2 | SLC22A16 |  |
| ZNF571 | SYVN1 |  |
|  | TMEM116 |  |
|  | TMPRSS15 |  |
|  | TPPP3 |  |

**Figure 3:**
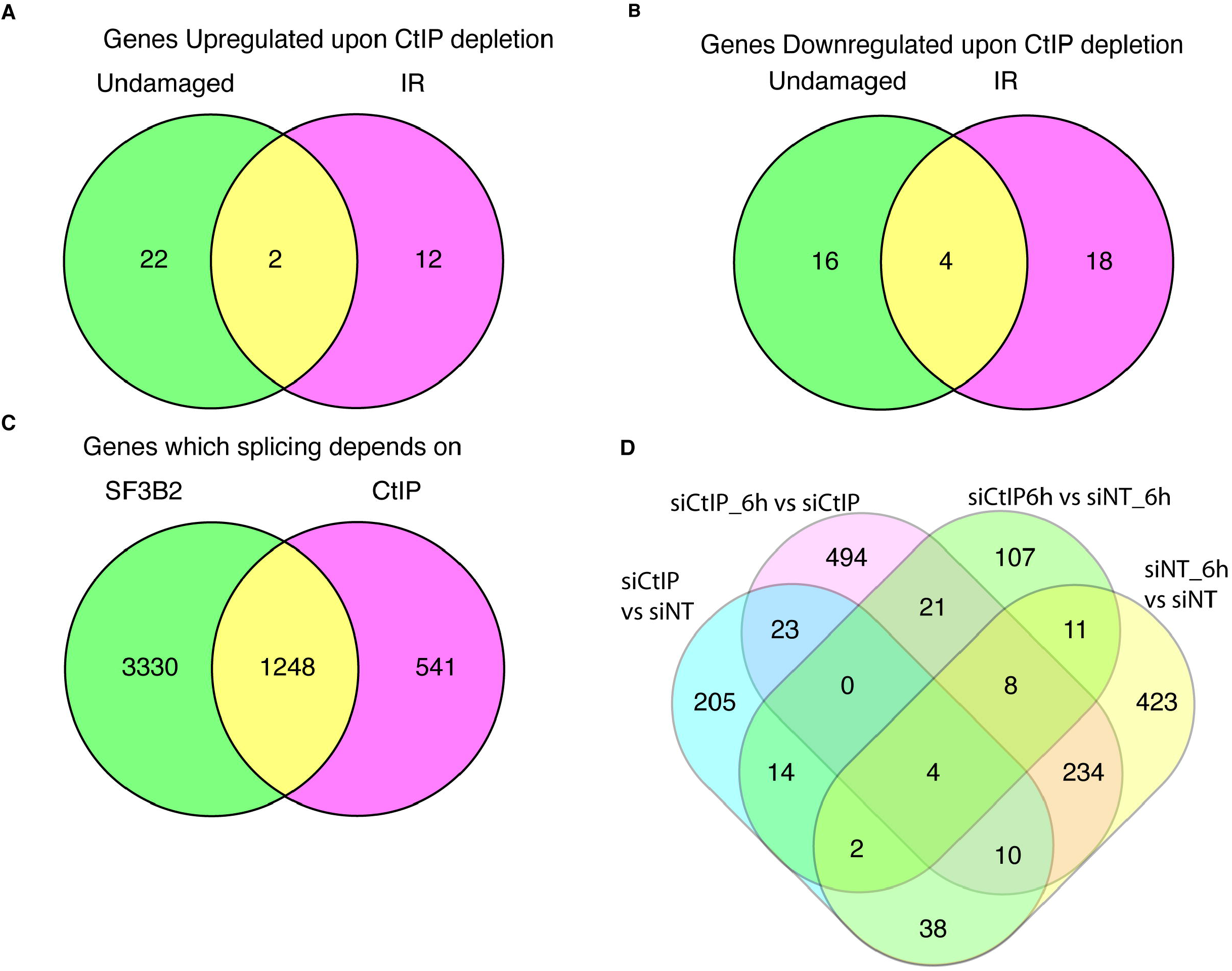
CtIP depletion affects gene expression and splicing of many genes. **A**, Distribution of the genes upregulated upon CtIP depletion regarding the exposure or not to DNA damage. Fold change (FC)>2, p-value<0.05. Gene expression was measured using the GeneChip HTA Array as described in the Materials and Methods section. The number of upregulated genes in undamaged cells (green), 6 h after exposure to irradiation (10 Gy; pink) or both (yellow) is shown in a Venn diagram. The actual list of genes can be found in Table 4. **B**, Same as A but for downregulated genes. The actual list of genes can be found in Table 5. **C**, the GeneChip HTA Array was used to look for genes that changed their splicing upon SF3B2 and/or CtIP depletion, as mentioned in the Materials and Methods section. The Venn digram represents the one that change when CtIP (pink), SF3B2 (green) or both are downregulated. Other details as in A. **D**, Differential splicing events bteween different conditions: undamaged cells depleted for CtIP (siCtIP) or transfected with a non-target siRNA (siNT); or RNA collected 6h after irradiation in cells depleted for CtIP (siCtIP_6h) or control cells (siNT_6h).

Additionally, we studied mRNA splicing in the same conditions mentioned above and, interestingly, we realized that the downregulation of CtIP rendered a strong effect on RNA processing of hundreds of genes. As shown in Table 7, columns 4 and 5, CtIP downregulation on its own changed the pattern of mRNA splicing compared to control conditions, even though this phenotype was more pronounced in unperturbed cells.

**Table 7:**
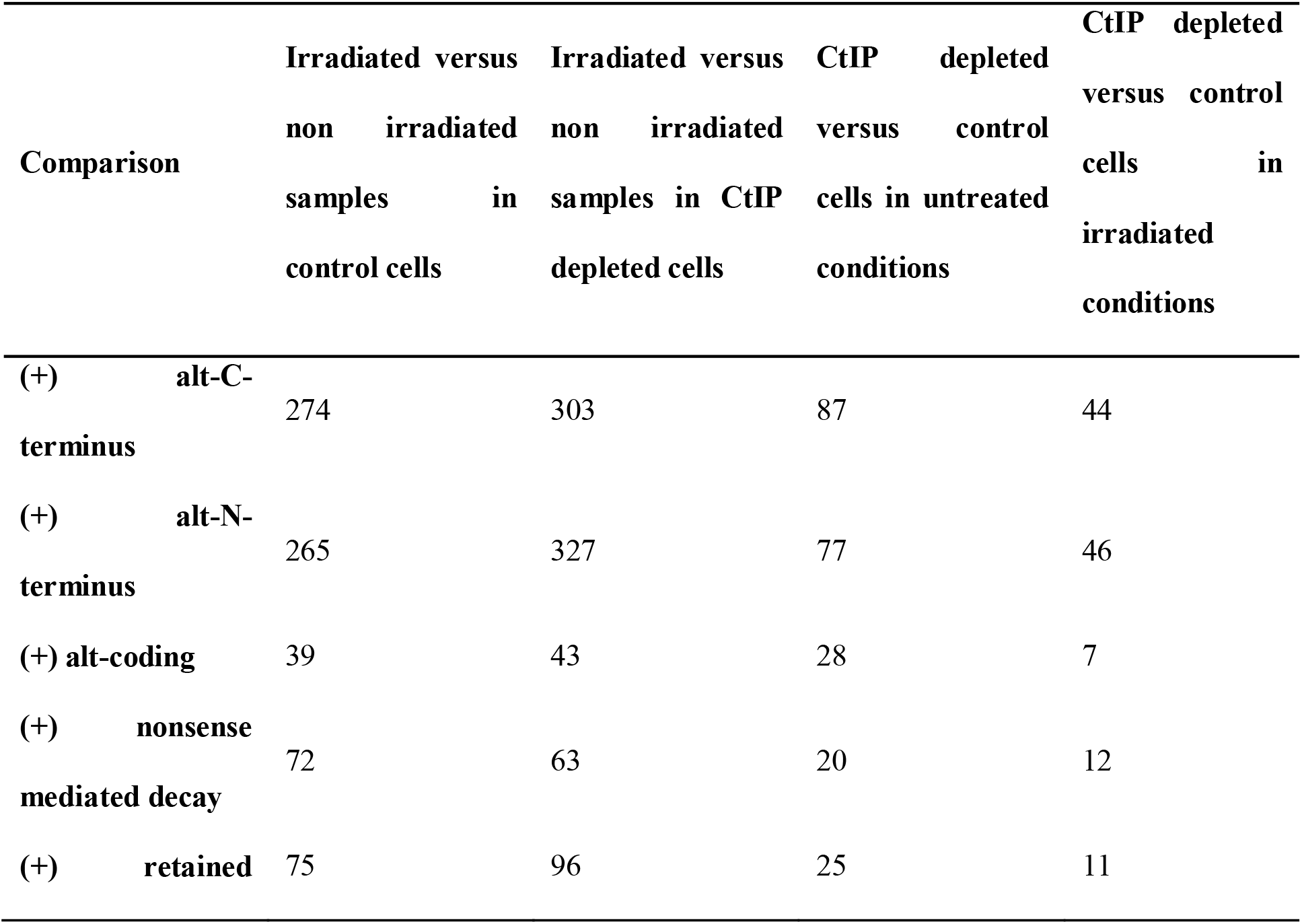
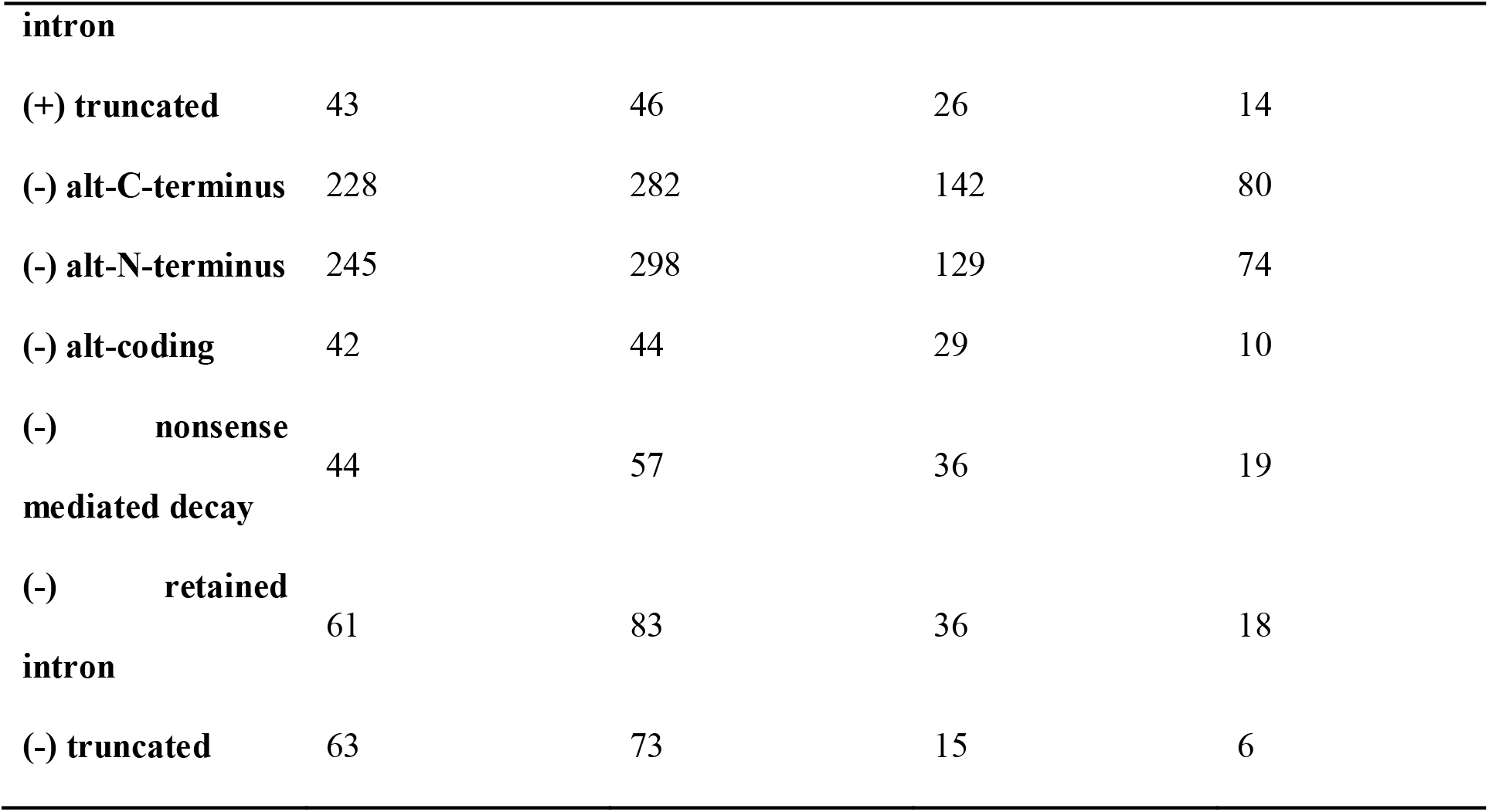
Specific splicing changes in response to IR and CtIP depletion. (+) represents gain and (-) loss of each specific event

As CtIP and SF3B2 physically interact, we wondered how many of the events that show a CtIP-dependent splicing did also rely on SF3B2 for that process, regardless of the exposure or not to DNA damage (Figure 3C). As expected, the number of splicing events that were dependent on SF3B2, a bona fide splicing factor, was higher than those that require CtIP. Indeed, only less than 30% of those SF3B2-dependent events were diminished upon CtIP depletion. But interestingly, around 70% of the CtIP-dependent splicing events were also affected by SF3B2. Thus, our results suggest that CtIP has a role in splicing, although less prominent than SF3B2. Moreover, most CtIP-dependent splicing events require also the SF3B complex, reinforcing the idea that CtIP might usually act with SF3B during splicing regulation, regardless to the fact that there are also a minority of CtIP-dependent but SF3B2-independent specific splicing events. On the contrary, SF3B can readily act on the splicing of most genes independently of CtIP.

Considering the role of CtIP in the response to DNA damage, we decided to simultaneously analyze the effect of CtIP depletion and irradiation on genome-wide alternative splicing. Thus, we carried out another analysis in which we compared all conditions in pairs: control cells without DNA damage (shNT) or exposed to IR (shNT6h) or depleted for CtIP in unperturbed (shCtIP) or damaged cells (shCtIP6h) (Figure 3D). Only 107 genes were altered due to CtIP absence in irradiated cells, whereas 205 did so in undamaged cells (Figure 3D; in green and blue respectively). The splicing of 494 genes was changed specifically in response to DNA damage exclusively in CtIP depleted cells (Figure 3D, in red). In a similar number of genes (423 genes; Figure 3D, yellow), mRNA splicing was modified in response to DNA damage only in cells that retained CtIP. We were particularly interested in those latter mRNA, as they represented DNA damage-dependent mRNA variants that are CtIP dependent. Hence, we reasoned that CtIP might mediate, directly or indirectly, such DNA damage-dependent alternative mRNA processing. Among them, we found that CtIP controls the DNA damage-dependent splicing of the helicase PIF1. Interestingly, SF3B2 also affects PIF1 splicing (Figure 2F). Strikingly, PIF1 and CtIP physically interact and together contribute to DNA end resection over specific DNA structures such as G-quadruplexes (Jimeno et al., 2018b). Thus, in order to study deeply the crosstalk between CtIP, DNA end resection and RNA splicing, we decided to focus on the altered splicing of PIF1.

### CtIP controls mRNA splicing of *PIF1*

PIF1 is a helicase with a 5’-3’ polarity. In humans there are only one PIF1 gene, but it was known to produce two well studied different transcripts (Supplementary Figure 1). A short transcript (2295nt) produces the longer protein isoform (707aa) called PIF1ß, which is located in the mitochondria. On the other hand, the longer transcript (2688nt) is translated into a smaller protein variant named PIF1α (641aa) that is localized in the nucleus (Futami et al., 2007). The difference between both proteins is the presence of a mitochondrial localization domain in PIF1ß, which also lacks the signal to translocate into the nucleus. Our array data showed additional splicing changes on the PIF1α backbone that were CtIP- and DNA damage-dependent (Figure 4A). We studied the inclusion of exon 3 (Figure 4A (I)), exon 4 (Figure 4A (II)), exon 9 (Figure 4A (III)) and exon 10 (Figure 4A (IV)). All these optional events are combinatorial and not mutually exclusive, so a mix of all the possible different species of mRNA coexist in all condition, regardless CtIP or DNA damage, but the array predicts changes on different conditions. We studied the abundance of the different alternative events by qPCR in cells depleted or not for CtIP with an siRNA and exposed or not to 10Gy of ionizing radiation (Figures 4B-G). The presence of an exon 2-3 junction was only slightly increased upon irradiation or CtIP depletion, but the changes were not significant (Figure 4C). A similar, but in this case significant, change was observed for exon 4 inclusion event (Figure 4D). Exon 8-9 junction presence was strongly increased in response to irradiation in control cells, and both its inclusion and DNA damage-accrue was completely dependent on CtIP presence (Figure 4E). Exon 10 inclusion event in the mRNA analysis rendered no statistically significant changes in any conditions (Figure 4F). These results suggested that the clearest CtIP-dependent alternative events in response to exogenous damage occur in the exon 4 inclusion and, more specially, in exon 8-9 junction of *PIF1* gene. Interestingly, *PIF1* mRNA splicing was also affected upon SF3B2 depletion (Figure 2F), although in this case we had analyzed the effect on the exon 9-10 junction as it was the prominent change observed in the array. In order to test if the 8-9 junction of PIF1 was also controlled by SF3B2 and if such effect was similar to CtIP, we repeated the qPCR experiments using the specific pair of oligos. Strikingly, the DNA damage-induced increase in the 8-9 junction of PIF1 was also controlled by SF3B2 (Figure 4G), but in a fashion that did not resemble the regulation by CtIP (Figure 4E) but the effect of SF3B2 on PIF1 exon 9-10 junction (Figure 2F). I. e., the depletion of SF3B2, instead of reducing this event like CtIP, generally increased it. However, both CtIP or SF3B2 knockdown abolished the DNA damage-dependent stimulation. Thus, we conclude that both SF3B2 and CtIP are required for the DNA-damage increase in the use of the exons 8 and 9 junction but have opposite effects in unchallenged conditions. To reinforce this idea, we repeated the analysis of the exon 8-9 junction in cells depleted of either factor upon stimulation of DNA damage with the topoisomerase I inhibitor camptothecin (CPT; Figure 4H). In agreement with our hypothesis, the results were similar to those obtained upon irradiation (compare Figure 4H with 4E and 4F).

**Figure 4:**
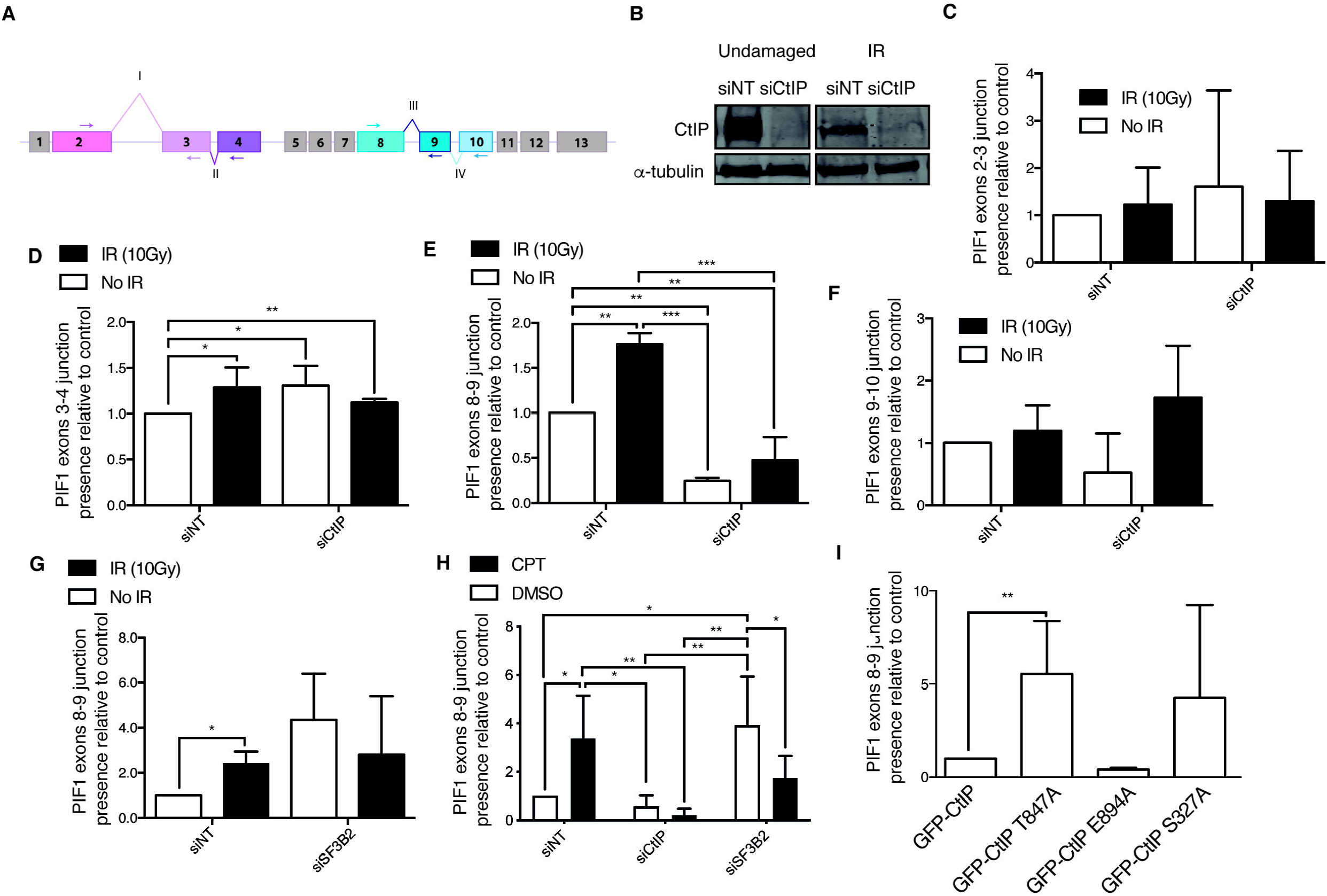
PIF1 splicing changes upon CtIP depletion. **A**, Schematic representation of PIF1 with the exons (boxes) and splicing events analyzed (roman numbers). **B**, Representative western blots showing the expression levels of CtIP upon depletion with a siRNA against CtIP or a control sequence (siNT) in cells exposed or not to 10 Gy of ionizing radiation. α-tubulin blot was used as loading control. **C**, Analysis of the exon 2 and exon 3 junction using quantitative PCR using primers located in exon 2 and 3 (see panel A) in cells depleted (siCtIP) or not (siNT) for CtIP, upon exposure to IR (black bars) or in unchallenged conditions (white bars). The data were normalized to the expression of a constitutive exon. The abundance of such event was normalized to the control cells in undamaged conditions. The average and standard deviation of three independent experiments is shown. Statistical significance was calculated using an ANOVA test. * p<0.05,** p<0.01 and *** p<0.005. **D**, same as B but for the inclusion of Exon 4. Primers in Exon 2 and 4 were used (see panel A). Other details as panel B. **E**, same as B but for the inclusion of Exon 9. Primers in Exon 8 and 9 were used (see panel A). Other details as panel B. **F**, same as B but for the inclusion of Exon 10. Primers in Exon 8 and 10 were used (see panel A). Other details as panel B. **G**, same as E but upon depletion of SF3B2 instead of CtIP. **H**, Same as E but upon depletion of either CtIP or SF3B2 as indicated and in cells exposed for 6h to camptothecin (CPT, black bars) or DMSO (with bars). **I**, Study of the exon 8-9 junction in cells depleted for endogenous CtIP and expressing the indicated mutants of CtIP. Other details as panel B.

In order to determine which activity of CtIP is involved in the splicing of *PIF1*, we carried out several qPCR to analyze the presence of the 8-9 junction on *PIF1* mRNA, but in cells bearing different mutated versions of CtIP (Figure 4I). We used GFP-CtIP as control, the resection defective CtIP-T847A, a CDK phosphorylation mutant (Huertas and Jackson, 2009) and CtIP-E894A, a sumoylation mutant (Soria-Bretones et al., 2017). Lastly, and considering that BRCA1 interacts with CtIP (Yu et al., 2006) and has been involved in damage-dependent alternative splicing (Savage et al., 2014), we analyzed the expression of *PIF1* exon 8-9 junction in the CtIP-S327A CDK phosphorylation mutant, that does not interact with BRCA1 but still resects albeit at a slower pace (Cruz-García et al., 2014). The data were normalized to the control GFP-CtIP, set as 1. Strikingly, both CDK defective phosphorylation mutants GFP-CtIP-T847A and GFP-CtIP-S327A caused a consistent increase in this splicing event, albeit only statistically significant in the T847A CtIP version (Figure 4I). This suggested CDK phosphorylation of CtIP inhibits the splicing of, at least, *PIF1* exon 8-9 junction. This is likely resection independent, as the E894A mutant, a sumoylation defective CtIP that is equally impaired in resection as the T847A mutant, did not share such phenotype (Figure 4G). Thus, specific posttranscriptional modifications seem also required for CtIP role in splicing. Specifically, CDK phosphorylation of CtIP blocks such role, suggesting that the splicing activity of CtIP happens mainly in G1.

### PIF1 splicing variants modulate DNA repair

Taking together these results, we decided to study the relevance of those CtIP- and DNA damage-dependent *PIF1* mRNA splicing events in DNA repair process in human cells. Hence, we created two different *PIF1* splicing variants cDNA constructs. First, a *PIF1* containing all exons (“total *PIF1*”; *tPIF1*), which correspond to the canonical PIF1α. In contrast, we also created a *PIF1* variant artificial cDNA that lacks both exon 4 and exon 9, the two exons which inclusion was more dependent on CtIP and DNA damage (Figures 4C and 4D). We named it “variant *PIF1*” (*vPIF1*). Importantly, such construct rendered a PIF1 that lacks part of the helicase domain of the protein. Both genes were expressed from pCDNA. In order to detect the expression of either variant in cells, we tagged both isoforms with GFP. An empty pCDNA plasmid was also used as control in our experiments. We transfected each plasmid (pCDNA, GFP-vPIF1 or GFP-tPIF1) into U2OS in order to study the effect of either isoform in human cells in response to DNA damage. Expression of the proteins coded by those variants is shown in Figure 5A.

**Figure 5:**
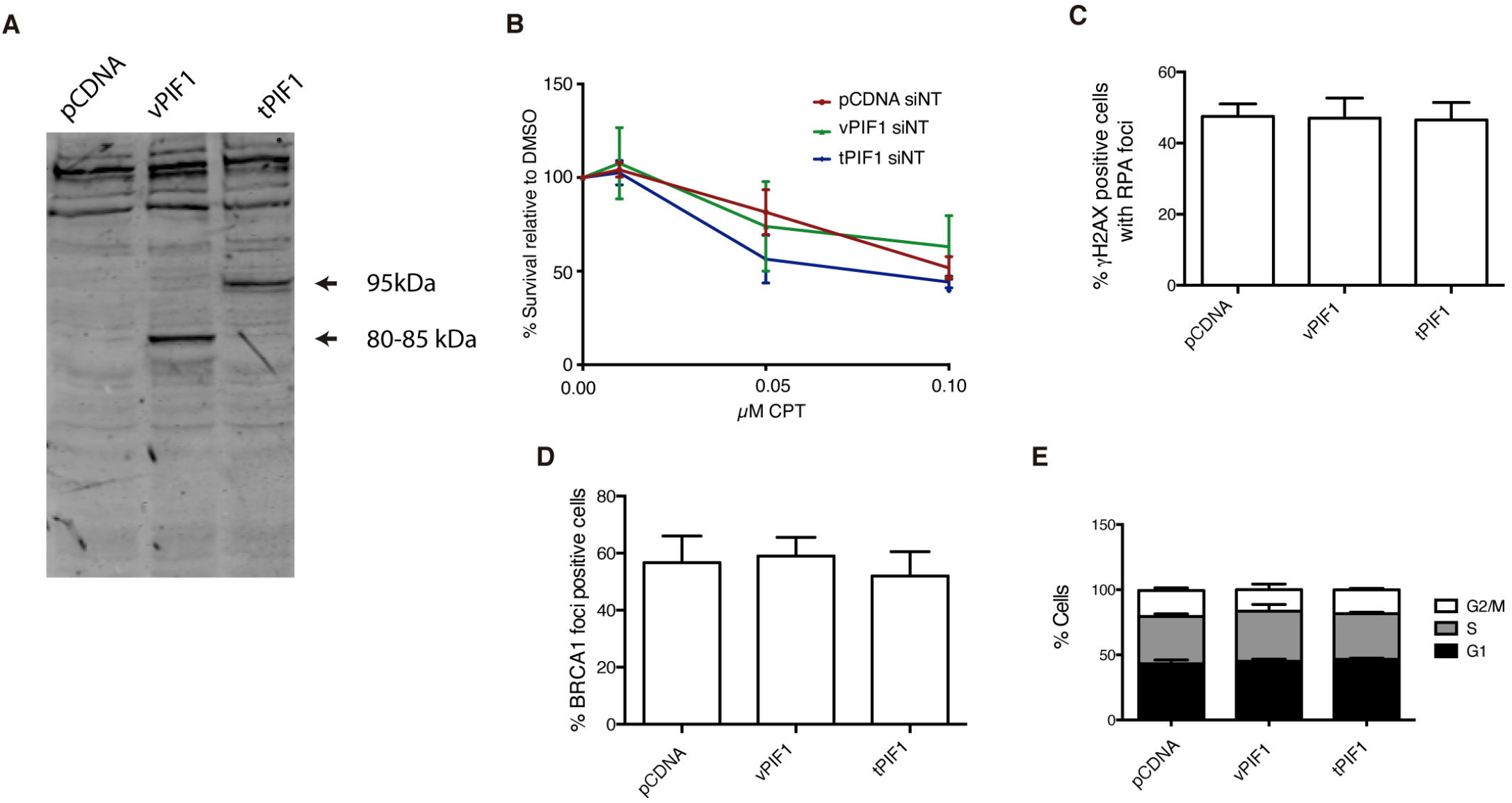
Expression of different PIF1 splicing variants. **A**, Western blot showing the abundance of different PIF1 isoforms in cells transfected with the empty pCDNA plasmid or pCDNA bearing the vPIF1 or tPIF1 splicing variants. **B**, Percentage of survival to different doses of camptothecin (CPT) in cells overexpressing the tPIF1 or vPIF1 isoforms relative to the DMSO treated control, as indicated. Cells transfected with the empty pCDNA vector were used as a control. The average and standard deviation of three independent experiments is shown. **C**, Percentage of cells positive for RPA foci upon exposure to 10 Gy of ionizing radiation. Cells expressed the indicated PIF1 variants. An empty pCDNA vector was used as a control. The average and standard deviation of three independent experiments is shown. No statistically significant differences were found using an ANOVA test. **D**, Same as A, but for BRCA1 foci. **E**, cell cycle analysis of cells transfected with the indicated plasmids. The percentage of cells in each cell cycle phase was analyzed as described in the Materials and Methods section. The average and standard deviation of three independent experiments is shown.

Considering that in all the conditions tested for the splicing analysis we could always observe a mixture of different splicing variant, including the canonical PIF1α we decided to leave the expression of the endogenous *PIF1* gene unperturbed, and combine it with the expression of the different PIF1 variants. First, we analyzed the ability of cells to survive to the DNA damaging agent camptothecin (CPT) when expressing the already spliced vPIF1 and tPIF1 constitutively. Strikingly, constant expression of the tPIF1 rendered cells sensitive to DNA damage when compared with cells expressing the empty plasmid (Figure 5B). This was not observed when the CtIP- and DNA damage-dependent spliced form vPIF1 was constitutively expressed (Figure 5B), suggesting that continuous expression of tPIF1 hampers DNA repair and, therefore, agreeing with the idea that a switch from tPIF1 to vPIF1 by DNA damage- and CtIP-induced alternative splicing ensures an adequate response.

Considering the fact that PIF1 and CtIP physically interact and cooperate in DNA end processing (Jimeno et al., 2018b), we decided to study DNA end resection in response to irradiation (10 Gy) in cells expressing either PIF1 construct. However, the overexpression of either isoform caused no effect in DNA resection or the recruitment of the resection and recombination factor BRCA1 (Figures 5C and D). As a control, we also analyzed the cell cycle in those cells to test whether such overexpression caused any change in the progress of cell cycle (Figure 5E). The lack of effect on RPA foci formation, an early event, and the timing of the observed splicing changes (6 h after irradiation) suggested that the transition from tPIF1 to vPIF expression might be more relevant for cell survival at later events of the DNA damage response and, therefore, it is separated from the resection role of PIF1.

### Early expression of vPIF1 delays DNA repair

In order to understand why changes in *PIF1* splicing to produce vPIF1 was only induced in response to DNA damage and that isoform was not constitutively expressed, we set to analyze the repair of DSBs at early time points in the presence of PIF1 isoforms. Interestingly, we observed that constitutive expression of vPIF1, albeit enhancing the long-term survival in response to DNA damage (Figure 5B) hampers or delays the recruitment of both NHEJ and HR proteins. Indeed, early after irradiation, the recruitment of the NHEJ factors 53BP1 and RIF1 was mildly impaired (Figure 6A and 6B). More strikingly, the recruitment of the essential HR factor RAD51 was severely impaired by constitutive expression of vPIF1 (Figure 6C). This was not observed when tPIF1 was overexpressed, confirming that this isoform does not block repair. Thus, our data suggest that a timely expression of different isoforms of PIF1 fine-tunes the response to DNA damage. Indeed, it seems that tPIF1 presence is permissive for early events, such the recruitment of 53BP1, RIF1 or RAD51, but in the long term is deleterious for cell survival in response to DSBs. vPIF1, on the contrary, increases the resistance to DNA damaging agents, despite the fact that expressed too early slows down DSB repair.

**Figure 6:**
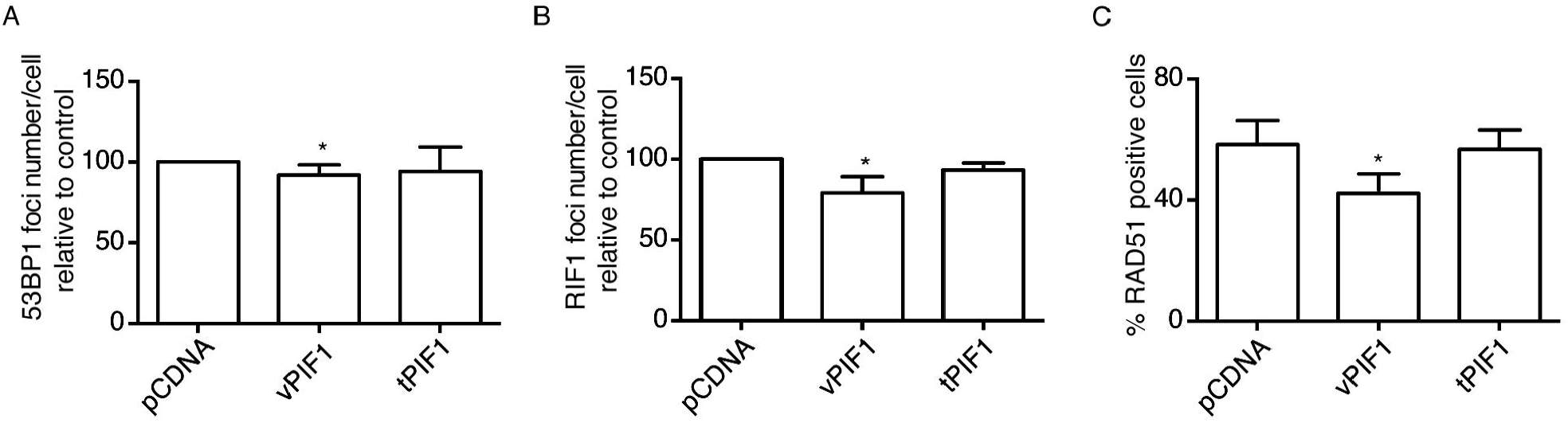
PIF1 splicing variants affect the recruitment of DDR factors. **A**, Average number of 53BP1 foci per cell upon exposure to 10Gy of IR in cells transfected with the indicated vectors relative to control cells. The average and standard deviation of three independent experiments is shown. Statistical significance was calculated using an ANOVA test. * p<0.05. **B**, same as A but for RIF1 foci. **C**, same as A but for RAD51 foci.

### Differential localization of PIF1 variants

As mentioned, there are two well-characterized PIF1 isoforms, the nuclear PIF1α and the mitochondrial PIF1β (Supplementary Figure 1). Although both our constructs, tPIF1 and vPIF1, are based on the nuclear form, PIF1α, we wondered whether vPIF1, which lacks part of the helicase domain, might show a different localization. To test it, we performed a cell fractionation assay. As shown Figure 7, the protein produced by the tPIF1 construct was mostly chromatin associated as expected (Figure 7A; green arrow): around 65% in the chromatin-bound and 35% in the cytoplasm (Figure 7B). On the contrary, the vPIF1 had the opposite distribution (Figure 7A; red arrow), with 65% of the protein in the cytoplasm and only 35% located in the chromatin fraction (Figure 7B). Those localizations did not change if cells were exposed to exogenous DNA damage (Data not shown).

**Figure 7:**
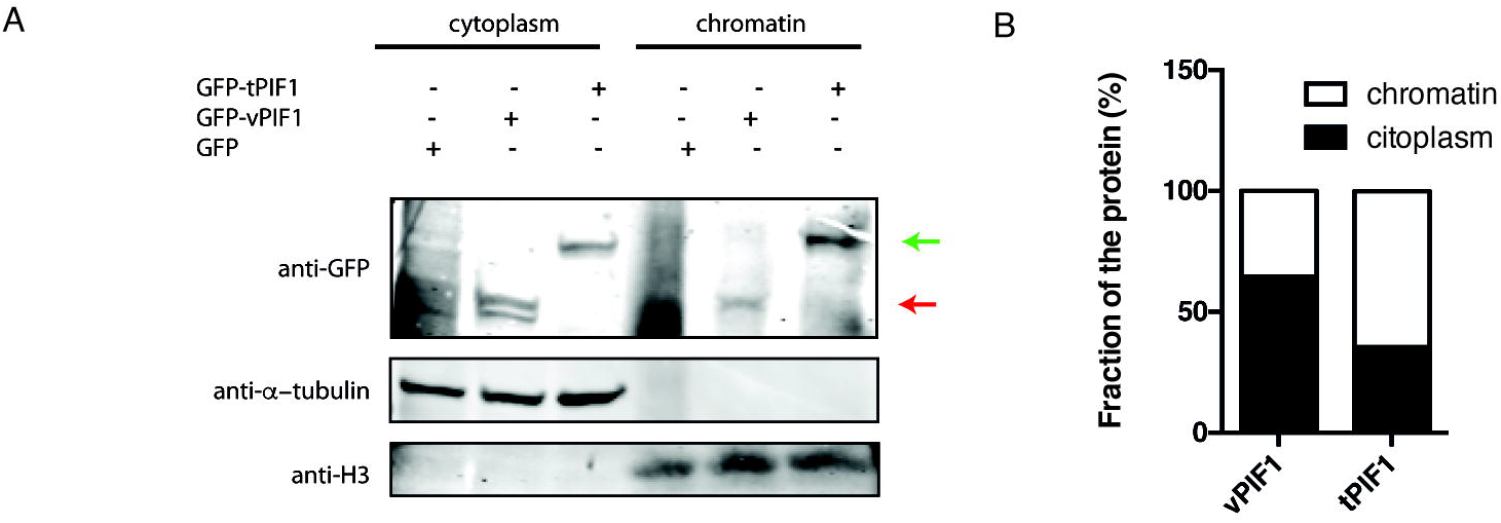
Cellular localization of PIF1 splicing variants. **A**, Protein samples from undamaged cells expressing the indicated PIF1 isoforms were fractionated as described in the Materials and Methods section. Cytoplasmic and chromatin fractions were resolved in SDS-PAGE and blotted with the indicated antibodies. vPIF1 is marked with a red arrow. tPIF1 is located with a green arrow. A representative experiment is shown. **B**, Western blots from A were quantified using an Odyssey Infrared Imaging System (LI-COR) and images ImageStudio software (LI-COR). The ration between the cytoplasmic and chromatin fraction of each PIF1 isoform is represented.

## DISCUSSION

In this work, we have discovered that not only the splicing complex SF3B but also the DNA repair factor CtIP controls the mRNA splicing of several genes, including proteins involved in DDR. Additionally, we have characterized the relevance of some splicing events on *PIF1* gene dependent on CtIP and DNA damage for the DDR and DNA repair.

Our data suggest that the SF3B complex controls the splicing and abundance of different mRNAs, both under unchallenged conditions and especially as a response to DNA damage, as a large set of genes have a differential splicing in SF3B2 depleted cells compared with control cells specifically upon exposure to ionizing irradiation. Interestingly, many of those genes are related RNA metabolism and protein modifications, known targets of the DNA damage response (Polo and Jackson, 2011). These DNA damage-dependent changes in mRNA metabolism agree with previous studies that reported large changes in mRNA expression (Gasch et al., 2001; Rieger et al., 2004). Additionally, several members of the SF3B complex have been identified by different genome-wide analysis as targets of the DDR (Beli et al., 2012; Bennetzen et al., 2010; Elia et al., 2015; Matsuoka et al., 2007) and DNA damage-dependent splicing events that are specific of certain splicing factors have been previously documented (Cloutier et al., 2018; Shkreta et al., 2016).. In SF3B case, we hypothesize that both aspects, gene expression and splicing changes, are indeed related to the splicing activity of the complex. Although regulation of transcription has been primarily associated for such alterations in gene expression, it has been increasingly clear that post-transcriptional modifications could indeed affect mRNA levels in response to DNA damage. Indeed, up to 50% of the changes on mRNA level in response to genotoxic agents could be attributed to mRNA turnover and not transcription (Boucas et al., 2012; Fan et al., 2002). Alternative splicing is known to affect mRNA stability in response to DNA damage by creating non-productive transcripts that are subjected to degradation by the non-sense mediated decay pathway (Barbier et al., 2007; Ip et al., 2011). In other cases, direct splicing-dependent gene expression repression or activation in response to DNA damage has been observed (Ip et al., 2011; Pleiss et al., 2007). Along those lines, we propose that alternative splicing controlled by the SF3B complex affects generally expression levels and the accumulation of alternative spliced mRNA in response to DNA damage. This global response will help the cells to fine-tune the response to broken DNA. Additionally, our splicing array data shows that SF3B2 affects also the splicing of key DDR factors, such as *BRCA1, RAD51, RIF1, DNA2* and *EXO1*. This might reflect a critical role of the SF3B complex in preparing the cells to the exposure to genotoxic agents.

Similarly, we show that CtIP affects the mRNA accumulation of hundreds of transcripts. This agrees, in principle, with the established role of CtIP in transcription (Koipally and Georgopoulos, 2002; Li et al., 1999; Liao et al., 2010; Liu and Lee, 2006; Liu et al., 2014; Sum et al., 2002; Xu et al., 2007). However, we also observe a prominent effect in splicing genome wide. Such role seems to require the functional interaction with the SF3B complex, as the majority of those events were altered also when SF3B2 was depleted. Splicing occurs cotranscriptionally and there is an intense crosstalk between transcription efficiency and splicing (Aslanzadeh et al., 2018; Braunschweig et al., 2014; Fong et al., 2014; Howe et al., 2003; Ip et al., 2011). Thus, transcription impairment could modify splicing efficiency and, on the contrary and as discussed for SF3B above, defective splicing might affect accumulation of mRNA that can be, erroneously, interpreted as transcriptional defects. So, in the case of CtIP is not so clear if those two roles, in transcription and splicing, are really two independent functions or simply both sides of the same coin. In any case, the regulation of the accumulation of different species of mRNA of many genes might explain why CtIP seems to play so many different roles in many processes. Indeed, there are evidences of other cases in which a RNA metabolism protein modulates its role in DDR through regulating the splicing of other factors involved in that process (Pederiva et al., 2016; Savage et al., 2014; Shkreta and Chabot, 2015). For example, PRMT5 regulates its effects on DNA repair by controlling the RNA splicing of several epigenetic regulators, especially the histone H4 acetyl-transferase TIP60 (KAT5) and the histone H4 methyltransferase SUV4-20H2 (KMT5C) (Hamard et al., 2018).

We propose that, for many phenotypes, and specially including DNA end resection, recombination and the DNA damage response, CtIP and SF3B probably participates at two different levels, directly, working as repair factors (Prados-Carvajal et al., 2018), but also indirectly as general transcription/splicing regulators. In that regard, they closely resemble its interactor BRCA1, which also seems to be involved at many different levels. Such double involvement of some RNA factors in DNA repair, which has simultaneous RNA-mediated and RNA-indirect roles in homologous recombination, was also observed by other authors (Anantha et al., 2013). This creates complex regulatory networks that are able to integrate multiple cellular signals and elicit sophisticated response that fine-tune the response.

Other proteins are likely involved in such complex networks, including the helicase PIF1. Interestingly, PIF1 and CtIP directly interacts and participates in resection of DSBs at atypical DNA structures such as G-quadruplexes (Jimeno et al., 2018b). But additionally, we have shown here that CtIP controls a DNA damage alternative splicing of PIF1 that modulates the response to DNA damaging agents. A proficient DDR seems to require a timely switch between the two forms described here, the tPIF1 and vPIF1. Whereas vPIF1 slightly impairs early events in DNA repair, its presence ensures a better survival to DNA damage. On the contrary, the presence of tPIF1 does not affect those early events, but if maintained in time compromise viability of cells exposed to camptothecin. Interestingly, the main difference between both forms is the change in cellular localization and the presence of an active helicase domain. Whereas tPIF1 maintains such activity, vPIF1 lacks exon 9 and, therefore, misses part of the active site. Thus, it is likely that tPIF1 participates on DNA end resection and DNA repair early on, working on a DNA substrate as a helicase. However, our data suggest that cells prefer to reduce this active PIF1 pool to ensure survival. One possibility is that the switch between both isoforms reduces the active pool of the helicase, both by sending the protein to the cytoplasm and by destroying the active site, to avoid interference of the helicase with very late steps of DNA repair or the DDR. Alternatively, it is formally possible that the vPIF1 plays some active role in the cytoplasm that facilitates survival. Considering that another isoform, PIF1βis essential for mitochondrial metabolism, we could not exclude this hypothetical function, although it will imply a role that does not require a functional helicase domain. In any case, PIF1 alternative splicing illustrates how CtIP might have additional effects in the response to DNA damage that has been, so far, overlooked. Strikingly, our data suggest that the regulation of PIF1 is modulated by the threonine 847 of CtIP, although not because resection is impaired of this mutant. This residue is a well stablished CDK site (Huertas, 2010; Polato et al., 2014). Interestingly, and albeit less clearly, CDK phosphorylation of CtIP S327 seems also involved. Hence, it is possible that upon DNA damage but only in cells in the G1 phase of the cell cycle, CtIP activates the expression of the isoform called vPIF1. This will separate the different roles of CtIP in a cell cycle dependent manner, with the DNA damage-induced splicing function mainly on G1 and the DNA end resection and homologous recombination exclusively in S and G2. Similar differential roles to maintain genome stability during the cell cycle have been reported for other factors before. For example, Rad^4TopBP1^ selectively activates checkpoint responses to DNA damage or replication perturbation depending on the cell cycle (Taricani and Wang, 2006).

In agreement with a tight relationship between BRCA1, CtIP and the SF3B complex, all three are intimately related to cancer appearance and, more specifically, with breast cancer incidence (Gokmen-Polar et al., 2019; Maguire et al., 2015; Paul and Paul, 2014; Soria-Bretones et al., 2013). In the case of CtIP and BRCA1, this connection with cancer has been mostly explained as a defective DNA repair. However, considering the involvement of CtIP, described here, and BRCA1 (Savage et al., 2014) in RNA splicing, maybe the cancer connection should be revisited on the light of those novel roles. Conversely, recent studies have detected recurrent mutations in components of the spliceosome in myelodysplastic syndromes (Pollyea et al., 2019; Shiozawa et al., 2018), renal cell carcinoma (Verma and Das, 2018; Yang et al., 2017), chronic lymphocytic leukaemia (Agrawal et al., 2017; Maleki et al., 2019), lung adenocarcinoma (Kim et al., 2018; Mao et al., 2019), breast cancer (Gokmen-Polar et al., 2019; Zhao et al., 2019) or pancreatic cancer (Tian et al., 2015; Zhou et al., 2017). Moreover, alterations in expression of splicing factors, including SF3B, can derive in various types of cancers (Alsafadi et al., 2016; Goswami et al., 2014; Maguire et al., 2015; Zheng et al., 2018). This probably reflects the importance of mRNA splice variants of several significant genes in apoptosis, metabolism, and angiogenesis (Grosso et al., 2008). However, another tantalizing possibility, not yet analysed in detail, is that some of those connections of SF3B with cancer might be a consequence of its more direct role in DNA repair. Furthermore, it might be possible to exploit the defective DNA repair and DDR in SF3B deficient cancer for therapeutic interventions. Thus, this crosstalk between DNA repair and the DDR and splicing might become in the future an important target for cancer treatment.

## MATERIALS AND METHODS

### Cell lines and growth conditions

U2OS human cell lines were grown in DMEM (Sigma-Aldrich) supplemented with 10% fetal bovine serum (Sigma-Aldrich), 2 mM L-glutamine (Sigma-Aldrich) and 100 units/ml penicillin and 100 µg/ml streptomycin (Sigma-Aldrich). For cells expressing GFP-tPIF1 and GFP-vPIF1, medium was supplemented with 0.5 mg/ml G418 (Sigma).

### shRNAs, siRNAs, plasmids and transfections

shRNAs and siRNA duplexes were obtained from Sigma-Aldrich, Dharmacon or Qiagen (Table 8) and were transfected using RNAiMax Lipofectamine Reagent Mix (Life Technologies), according to the manufacturer’s instructions. Plasmid transfection of U2OS cells with PIF1 variants was carried out using FuGENE 6 Transfection Reagent (Promega) according to the manufacturer’s protocol.

**Table 8:**
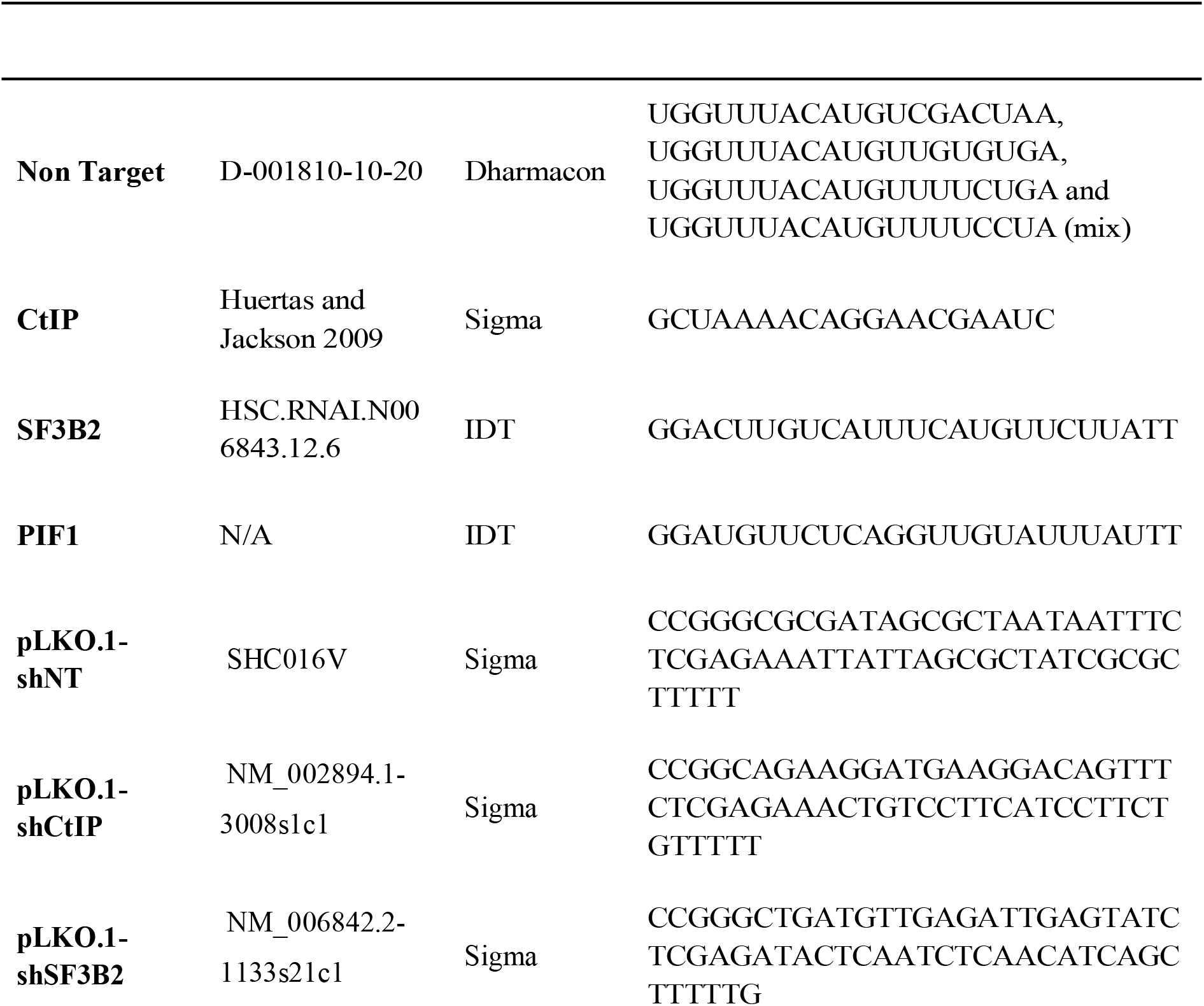
shRNAs and siRNAs used.

To generate the lentivirus harboring the shRNA, HEK293T cells were transfected with the plasmids containing the specific shRNA (pLKO.1), p8.91 and pVSVG in a ratio 3:2:1. The DNA mix was prepared in a volume of 1 ml containing 250 mM CaCl2, and was added dropwise while bubbling into 1 ml of 2x HEPES buffered saline (Sigma). The mix was incubated 30 minutes at room temperature and added dropwise to the cells while carefully rocking the plate. Two days after transfection, medium was collected and filtered using 0.45 μm polyvinylidene difluoride (PVDF) filters (Millipore) and 8 μg/ml polybrene (Sigma,) was added. For transduction, cells were seeded, and 24 hours later medium with lentiviruses was thawed and added to the plates. The medium was replaced 8 hours later to remove viral residues and polybrene. Knockdowns were validated by western blot 48h after transduction.

### Cell fractionation

U2OS cells stably expressing the different PIF1 isoforms were subjected to nuclear/cytoplasm fractionation to analyze the distribution of both PIF1 variants in the different cellular compartments basically following the protocol described by Philpott and colleagues (Gillotin et al., 2018). Briefly, after washing the cells once with room temperature PBS, they were resuspended into 1 ml PBS and then pe pelleted at 1,200 rpm at 4 °C for 3 min. Pellets were covered with 5 volumes of ice-cold E1 buffer (50 mM Hepes-KOH pH 7.5, 140 mM NaCl, 1 mM EDTA pH8.0, 10% glycerol, 0.5% NP-40, 0.25% triton X-100, 1 mM DTT) complemented with 1 × protease inhibitor cocktail and gently pipetted up and down (5 times). After centrifugation at 3,400 rpm at 4 °C for 2 min, the supernatant was collected (cytoplasmic fraction). Then the pellet was washed in the same volume of E1 buffer. The pellet was resuspended in the same volume of E1 and incubated for 10 min on ice. Upon centrifugation at 3,400 rpm at 4 °C for 2 min, the supernatant was discarded. The pellet was resuspended by gentle pipetting in 2 volumes of ice-cold E2 buffer (10 mM Tris-HCl pH8.0, 200 mM NaCl, 1 mM EDTA pH8.0, 0.5 mM EGTA pH8.0) complemented with 1 × protease inhibitor cocktail and then centrifuged at 3,400 rpm at 4 °C for 2 min. The supernatant was collected in a fresh tube (nucleus fraction). The pellet was washed in the same volume of E2, resuspended in the same volume of E2 and incubated for 10 min on ice. Upon centrifugation at 3,400 rpm at 4 °C for 2 min, the supernatant was discarded and the pellet was resuspended in ice-cold E3 buffer (500 mM Tris-HCl, 500 mM NaCl) complemented with 1 × protease inhibitor cocktail. The solution (chromatin fraction) was sonicated to solubilize the proteins. All fractions were then centrifuged at 13,000 rpm at 4 °C for 10 min to clarify the solution.

### SDS-PAGE and Western blot analysis

Protein extracts were prepared in 2× Laemmli buffer (4% SDS, 20% glycerol, 125 mM Tris-HCl, pH 6.8) and passed 10 times through a 0.5 mm needle–mounted syringe to reduce viscosity. Proteins were resolved by SDS-PAGE and transferred to low fluorescence PVDF membranes (Immobilon-FL, Millipore). Membranes were blocked with Odyssey Blocking Buffer (LI-COR) and blotted with the appropriate primary antibody (Table 9) and infra-red dyed secondary antibodies (LI-COR) (Table 10). Antibodies were prepared in Blocking Buffer supplemented with 0.1% Tween-20. Membranes were air-dried in the dark and scanned in an Odyssey Infrared Imaging System (LI-COR), and images were analyzed with ImageStudio software (LI-COR).

**Table 9:** Primary antibodies used in this study. WB, western blotting. IF, immunofluorescence.

| Target protein | Source | Supplier/Reference | Application | Dilution |
| --- | --- | --- | --- | --- |
| GFP | Rabbit | Santa Cruz (sc-8334) | WB | 1:1000 |
| α-tubulin | Mouse | Sigma (T9026) | WB | 1:50000 |
| β-actin | Rabbit | Abcam (ab8227) | WB | 1:50000 |
| RPA32 | Mouse | Abcam (ab2175) | IF | 1:500 |
| γH2AX | Rabbit | Cell Signaling (2577L) | IF | 1:500 |
| RAD51 | Mouse | Abcam; ab213 | IF | 1:1000 |
| BRCA1 | Mouse | Santa Cruz (sc-6954) | WB, IF | 1:1000, 1:200 |
| 53BP1 | Rabbit | NB100-304, Novus | WB, IF | 1:1000, 1:500 |
| RIF1 | Mouse | Bethyl Laboratories<br>(A300-568A) | WB, IF | 1:500, 1:200 |
| CtIP | Mouse | R. Baer (14.1) | WB | 1:500 |
| SF3B2 | Rabbit | Proteintech (10919-1-AP) | WB | 1:1000 |
| PIF1 | Mouse | Santa cruz (sc-48377) | WB | 1:500 |

**Table 10:** Secondary antibodies used in this study. WB, western blotting. IF, immunofluorescence

| Antibody | Supplier/Reference | Application | Dilution |
| --- | --- | --- | --- |
| IRDye 680RD goat anti-mouse IgG (H+L) | LI-COR (926-68070) | WB | 1:10000 |
| IRDye 800CW goat anti-rabbit IgG (H+L) | LI-COR (926-32211) | WB | 1:10000 |
| Alexa Fluor 594 goat anti-mouse | Invitrogen (A11032) | IF | 1:1000 |
| Alexa Fluor 488 goat anti-rabbit | Invitrogen (A11034) | IF | 1:1000 |

### Immunofluorescence and microscopy

For RPA foci visualization, U2OS cells knocked-down for different proteins were seeded on coverslips. At 1 h after either irradiation (10 Gy), coverslips were washed once with PBS followed by treatment with Pre-extraction Buffer (25 mM Tris-HCl, pH 7.5, 50 mM NaCl, 1 mM EDTA, 3 mM MgCl2, 300 mM sucrose and 0.2% Triton X-100) for 5 min on ice. To visualize RAD51 foci, cells were cultured for 3 h after irradiation. For visualize BRCA1 foci, Pre-extraction Buffer 2 (10mM PIPES pH 6.8, 300 mM sucrose, 50mM Nacl, 3mM EDTA, 25X proteinase inhibitor and 0.5% Triton X-100) was used. Cells were fixed with 4% paraformaldehyde (w/v) in PBS for 15 min. Following two washes with PBS, cells were blocked for 1 h with 5% FBS in PBS, co-stained with the appropriate primary antibodies (Table 9) in blocking solution overnight at 4°C or for 2 h at room temperature, washed again with PBS and then co-immunostained with the appropriate secondary antibodies for 1 h (Table 10) in Blocking Buffer. After washing with PBS and drying with ethanol 70% and 100% washes, coverslips were mounted into glass slides using Vectashield mounting medium with DAPI (Vector Laboratories). RPA foci immunofluorescences were analyzed using a Leica Fluorescence microscope.

For 53BP1 visualization, U2OS cells were seeded and transfected as previous described. Once collected, cells were fixed with methanol (VWR) for 10 min on ice, followed by treatment with acetone (Sigma) for 30 sec on ice. For RIF1 foci visualization, cells were fixed with 4% PFA for 15 min, washed twice with 1× PBS and then permeabilized for 15 min with 0.25% Triton diluted in 1× PBS. Samples were immunostained as described above with the appropriate primary and secondary antibodies (Tables 9 and 10). Images obtained with a Leica Fluorescence microscope were then analyzed using Metamorph to count the number of foci per cell.

### Cell cycle analysis

Cells were fixed with cold 70% ethanol overnight, incubated with 250 μg/ml RNase A (Sigma) and 10 μg/ml propidium iodide (Fluka) at 37°C for 30 min. For each replicate, 10.000 cells were analyzed with a FACSCalibur (BD). Cell cycle distribution data were further analyzed using ModFit LT 3.0 software (Verity Software House Inc).

### RNA extraction, reverse transcription and quantitative PCR

RNA extracts were obtained from cells using NZY Total RNA Isolation kit (Nzytech) according to manufacturer’s instructions. To obtain complementary DNA (cDNA), 1 μ g RNA was subjected to RQ1 DNase treatment (Promega) prior to reverse transcription reaction using Maxima H Minus First Strand cDNA Synthesis kit (Thermo Scientific) according to manufacturer’s instructions. Quantitative PCR from cDNA was performed to check siRNA-mediated knock-down of several proteins. For this, iTaq Universal SYBR Green Supermix (Bio-Rad) was used following manufacturer’s instructions. DNA primers used for qPCR are listed in Table 4. Q-PCR was performed in an Applied Biosystem 7500 FAST Real-Time PCR system. The comparative threshold cycle (Ct) method was used to determine relative transcripts levels (Bulletin 5279, Real-Time PCR Applications Guide, Bio-Rad), using *β*-actin expression as internal control. Expression levels relative to *β*-actin were determined with the formula 2^-ΔΔct^ (Livak and Schmittgen, 2001). To analyze *PIF1* exon junctions, the data were also normalized to an exon constitutively expressed in the cell.

### Microarray

To analyse RNA splicing genome wide from cells depleted or not for SF3B2 or CtIP in damaged (10 Gy) and untreated conditions the splicing microarray Transcriptome Arrays HTA & MTA (Affymetrix) was used as previously described (Jimeno-González et al., 2015). This array design contains >6.0 million distinct probes covering coding and non-coding transcripts. 70% of the probes cover exons for coding transcripts, and 30% of probes on the array cover exon-exon splice junctions and non-coding transcripts. The array contains approximately ten probes per exon and four probes per exon-exon splice junction. RNA was obtained using an RNA isolation kit, as explaineed in section above, in triplicates. The purity and quality of isolated RNA were assessed by RNA 6000 Nano assay on a 2100 Bioanalyzer (Agilent Technologies). Total RNA from each sample (250 ng) was used to generate amplified and biotinylated sense-strand cDNA from the entire expressed genome according to the GeneChip WT PLUS Reagent Kit User Manual (P/N 703174 Rev 1; Affymetrix Inc.). GeneChip HTA Arrays were hybridized for 16 h in a 45 °C oven, rotated at 60 rpm. According to the GeneChip Expression Wash, Stain and Scan Manual (PN 702731 Rev 3; Affymetrix Inc.), the arrays were washed and stained using the Fluidics Station 450 and finally were scanned using the GeneChip Scanner 3000 7G. Raw data were extracted from the scanned images and analyzed with the Affymetrix Command Console 2.0 Software. The raw array data were preprocessed and normalized using the Robust Multichip Average method. Data were processed further using Transcriptome Analysis Console (TAC) Software from Affymetrix, which performs a gene-level analysis or an alternative splicing analysis. For gene-level analysis, gene expression changes with P < 0.05 (ANOVA) and a linear fold change| ≥2 were considered significant. For alternative splicing analysis, the splicing index (SI) was calculated as SI=Log2[(Probe−set1Doxintensity/Gene1Doxintensity)/(Probe−set1ControlIntensity/G ene1ControlIntensity)]. Splicing changes were considered significant when the |splicing index| was ≥2 and p < 0.05 (ANOVA).

Array data also were analyzed using AltAnalyze software (Emig et al., 2010) version 2.0 with core probe set filtering using DABG (detected above background; P value cutoff of 0.05) and microarray analysis of differential splicing (MiDAS exon analysis parameters; P value cutoff of 0.05). The signs of the splicing index values obtained from AltAnalyze were changed to use the same splicing index definition throughout the manuscript. AltAnalyze incorporates a library of splicing annotations from UCSC KnownAlt database. GeneChip HTA 2.0 arrays include probes to detect 245.349 different transcript variants, supported by a variable number of ESTs in the databases. Intronic regions often are represented in the ESTs databases because of the retrotranscription of unspliced pre-mRNAs, aberrant splicing forms, and spurious firing of cryptic promoters inside introns, with functional or nonfunctional implications. For that reason, GeneChip HTA 2.0 arrays allow the identification of an intron-retention event (media.affymetrix.com/support/technical/datasheets/hta_array_2_0_datasheet.pdf).

### Statistical analysis

Statistical significance was determined with a Student’s *t*-test or ANOVA as indicated using PRISM software (Graphpad Software Inc.). Statistically significant differences were labelled with one, two or three asterisks for *P* < 0.05, *P* < 0.01 or *P* < 0.001, respectively.

## ACKNOWLEDGEMENTS

We wish to thank Jose Carlos Reyes for critical reading of the manuscript. This work was financed by an R+D+I grant from the Spanish Ministry of Economy and Competitivity (SAF2016-74855-P) and by the European Union Regional Funds (FEDER). RP-C was funded with an FPU fellowship from the Spanish Ministry of Education, and GR-R was supported by the Regional Government of Andalucía (Junta de Andalucía) with a contract of the program “Garantía juvenil en la Universidad de Sevilla”. CABIMER is supported by the regional government of Andalucía (Junta de Andalucía).

