## Supplementary figures and images for "CtIP -mediated alternative mRNA splicing finetunes the DNA damage response"

### Supplementary Figure 1

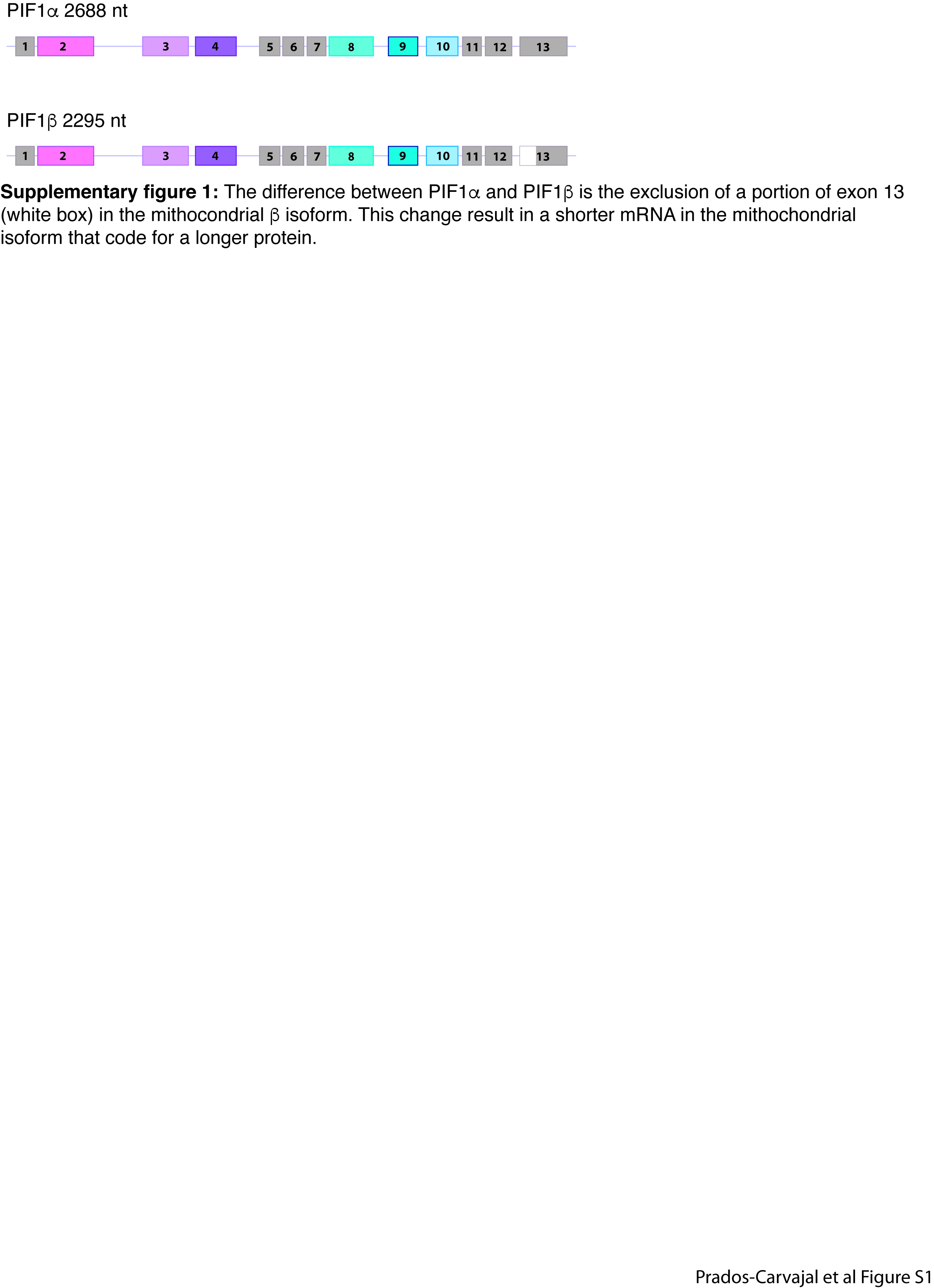
